# Potent activation of NAD^+^-dependent deacetylase Sirt7 by nucleosome binding

**DOI:** 10.1101/2022.05.11.491540

**Authors:** Vyacheslav I. Kuznetsov, Wallace H. Liu, Mark A. Klein, John M. Denu

## Abstract

Sirtuin-7 (Sirt7) is a nuclear NAD^+^-dependent deacetylase with a broad spectrum of biological functions. Sirt7 overexpression is linked to several pathological states and enhances anticancer drug resistance, making the enzyme a promising target for the development of novel therapeutics. Despite a plethora of reported *in vivo* functions the biochemical characterization of recombinant Sirt7 remains inadequate for the development of novel drug candidates. Here, we conduct an extensive biochemical analysis of Sirt7 using newly developed binding and kinetic assays to reveal that the enzyme preferentially interacts with and is activated by nucleosomes. Sirt7 activation by nucleic acids alone is effective towards long-chain acylated hydrophobic substrates while only nucleosome binding leads to 10^5^-fold activation of deacetylase activity. Using endogenous chromatin and recombinant acetylated nucleosomes, we reveal that Sirt7 is one of the most efficient deacetylases in the sirtuin family and that its catalytic activity is limited by the rate of dissociation from deacetylated nucleosomes.

---

The family of mammalian sirtuins comprises seven enzymes (Sirt1-7) that differ in subcellular localization, substrate specificity, interaction partners, and the level of intrinsic enzymatic activity.^1,2^ The enzymes can function by deacylating a broad range of proteins in a NAD^+^-dependent manner producing *O*-acetyl(acyl)-ADP ribose, nicotinamide, and modification-free target.^3^ Dependence of sirtuins’ activity on intracellular NAD^+^ level allows these enzymes to function as metabolic sensors that link NAD^+^ availability to specific intracellular protein deacetylation.^4^

Sirt7 is a 400 amino acid protein within the Class III deacetylase family of enzymes and features a C-terminal nuclear localization sequence (NLS) that shuttles Sirt7 into the nucleus.^5^ Sirt7 is further sub-localized to the nucleolar compartment likely due to the highly cationic arginine-rich protein surface.^6^ Sirt7 is expressed in multiple organs and tissues including skeletal muscle, heart, and brain with the highest level of expression detected in the spleen, liver, and testis.^7^ Sirt7 knockout in mouse model affects lifespan and aging, lipid^8,9^ and mitochondrial metabolism,^10^ development of cardiac disease.^11^ The overexpression of Sirt7 is strongly correlated with cellular proliferation and various types of aggressive cancer.^12–14^ Sirt7 increases survivability, drug, and radio-resistance of cancer cells,^15,16^ and together with PARP the enzyme moderates the therapy-induced cellular DNA damage.^17,18^ The diverse functional roles of Sirt7 under normal and pathological conditions make this enzyme a promising candidate for the drug development process.

Despite numerous biological studies, the biochemical aspects of Sirt7 activity and regulation remain poorly understood. Unlike other sirtuins, the recombinant apoSirt7 was found to be completely inactive, but when activated by nucleic acids the enzyme displayed a very strong preference for hydrophobic H3K18 acylated substrates while showing weak deacetylation activity.^19^ These *in vitro* observations are incongruous with numerous biological studies that demonstrate the ability of Sirt7 to effectively deacetylate multiple targets *in vivo*.^20^The apparent discrepancy could be explained by the existence of an unknown intracellular modulating mechanism that controls Sirt7 deacetylation activity and biological function *in vivo*. Understanding these fundamental factors is important for the prospective development of potent and selective drug candidates that must be screened against the active biologically relevant form of Sirt7.

In this work, we conducted an extensive *in vitro* evaluation of Sirt7 and analyzed the important interactions that control its short- and long-chain deacylation activity. For this purpose, we developed a novel high-yielding *E. coli* expression and purification strategy for Sirt7 and designed several innovative assays to analyze the binding affinity and activating potential of free nucleic acids (DNA, tRNA) and nucleosomes. We show that Sirt7 preferentially binds to nucleosomes resulting in profound activation of site-specific deacetylation activity. The substantial Sirt7 activation by DNA or tRNA is only observed with hydrophobic long-chain acylated substrates, while the deacetylation efficiency of these Sirt7-DNA and Sirt7-tRNA complexes remains very low. Finally, using acetylated recombinant nucleosomes as well as native oligo-nucleosomes isolated from deacetylase-inhibitor treated mammalian cells, we show that Sirt7 is a highly effective and site-specific chromatin deacetylase. Our results reveal how recombinant apoSirt7 deacetylation activity can be activated by site-specific interaction with nucleosomes that explain the existing functional discrepancies between biochemical (*in vitro*) and biological (*in vivo*) enzymatic behavior. Understanding how SIRT7 deacetylation function is regulated in the cellular contexts is crucial for the prospective development of specific pharmacological modulators against the active and biologically relevant Sirt7-nucleosome complex. In addition, the newly developed and optimized biochemical techniques can be directly applied for Sirt7-specific drug screening or more broadly used to study other nucleosome-binding proteins.

## Results and Discussion

### Sirt7 interacts with nucleic acids and nucleosomes

*In vivo* biological studies of Sirt7 had shown the ability of the enzyme to deacetylate several cellular targets^20^ including H3K18Ac^21^ chromatin mark. *In vitro*, the recombinant apoSirt7 was reported completely inactive, but activatable towards long-chain acylated peptides by complexation with DNA^19^ and to a greater extent with tRNA^22^ modulators. Low *in vitro* deacetylation activity of Sirt7, even with nucleic acids, suggested the existence of a more potent, but previously unknown Sirt7 activating mechanism. Considering the nuclear localization and previous chromatin deacetylation reports^21^ we tested nucleosomes as potential Sirt7 activators. To understand the enzymatic behavior of Sirt7 in the presence of various modulators (nucleic acids and nucleosomes), we first analyzed the stoichiometry and the complex speciation using electrophoretic mobility shift assay (EMSA). We chose tRNA as the representative RNA, as this form of RNA was previously reported to have the highest activating potential compared to other nucleic acids (DNA, rRNA).^22^ To investigate Sirt7-DNA interactions, we chose a highly purified 146bp Widom DNA that was also used to assemble recombinant human nucleosomes.^23^ Various Sirt7/DNA, Sirt7/tRNA, or Sirt7/nucleosome ratios were prepared, and the formed species were analyzed using EMSA. The resulting complexes were concurrently assessed for enzymatic activity towards a histone H3-derived synthetic peptide substrate (H3K18-Deca) using an improved HPLC assay (Figure 1, a-c). With 146bp double-stranded DNA as the interacting macromolecule (Figure 1, a), Sirt7 forms a complex mixture of distinct nucleoprotein species. The appearance of a family of sharp and proportionally spaced bands revealed by EMSA (Figure 1, a) is consistent with independent and tight interaction of multiple Sirt7 molecules with one equivalent of 146bp DNA. The saturated nucleoprotein complex forms at approximately 7-8 Sirt7 molecules per 146 bps with an average of 18-20 bps of DNA per enzyme equivalent. Sirt7 DNA interaction is coincident with an increase of long-chain H3K18Deca deacylation activity that reaches saturation when the Sirt7/DNA ratio approaches 6x-8x. At higher ratios, a decrease in activity is observed which coincided with a loss of distinct bands and the appearance of high molecular weight aggregates in the EMSA gel. Together, these data suggest that DNA binds and enhances Sirt7 activity toward the long-chain acylated H3K18Deca peptide substrate, but at high Sirt7/DNA ratio, the activating effect is diminished by the extensive aggregation.

**Figure 1.**
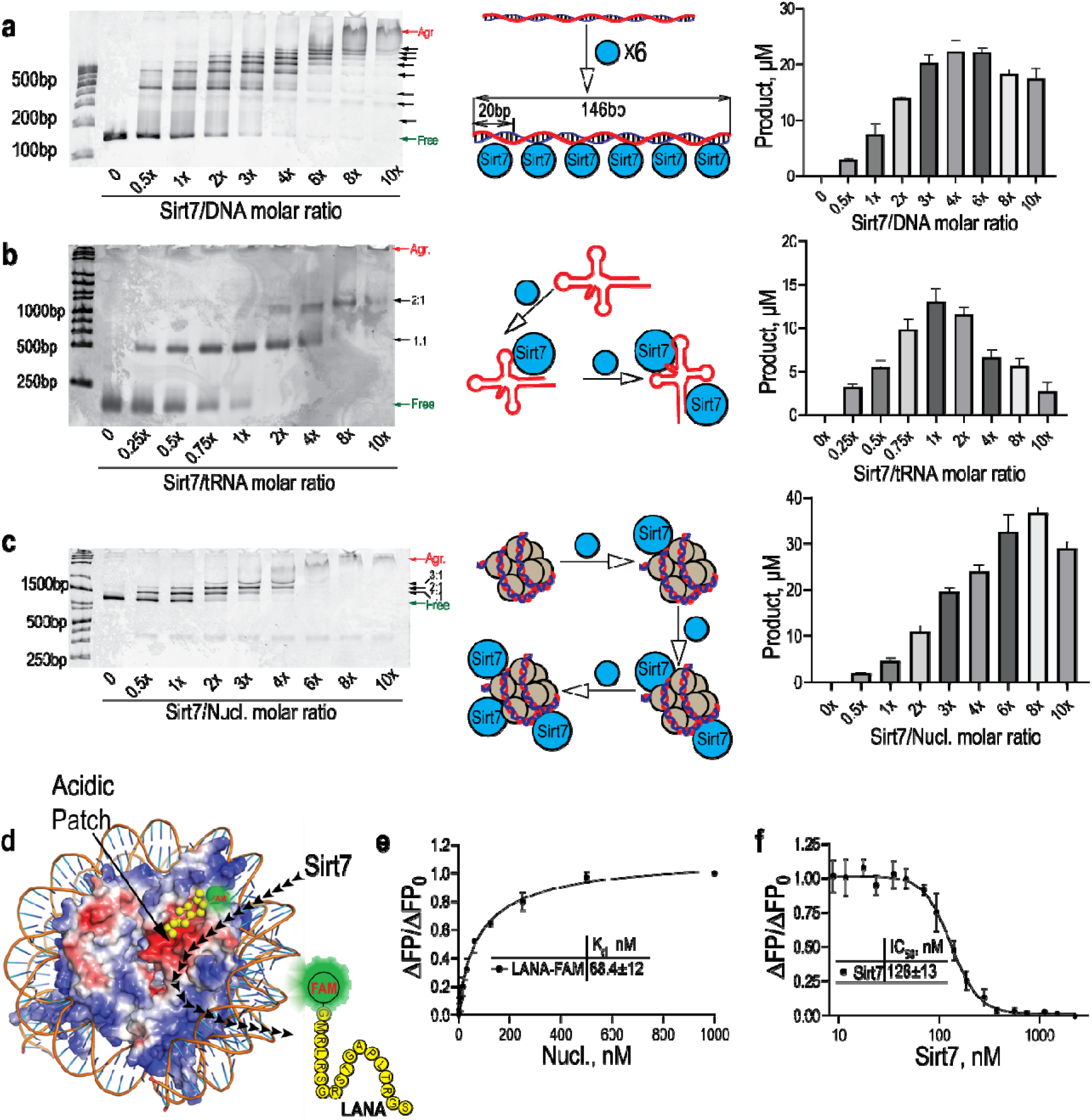
Analysis of Sirt7 interaction with DNA, tRNA, and nucleosomes. N tive EMSA gel analysis of Sirt7 interaction with Widom 601 DNA (a), tRNA (b), and nucleosome core particles (c). Sirt7 was titrated into 146bp DNA (a), tRNA (b), or nucleosomes (c), andformed species were identified by native EMSA. Detected complexes are marked arrows. The proposed interactional model and total enzymatic H3K18Deca deacyltion activity for each EMSA-analyzed Sirt7/macromolecule ratio is presented. d. Design of the LANA-FAM/Sirt7 competition experiment; e. FP analysis of LANA-FAM interaction nucleosomes. f. FP analysis of LANA-FAM displacement by Sirt7.

Unlike the 146 bp linear DNA that forms a poly-substituted nucleoprotein complex with Sirt7, the average 70-90 bp tRNA interaction with Sirt7 is more specific and stoichiometrically constrained. The gradual addition of Sirt7 to tRNA resulted in the sequential formation of mono- and di-substituted complexes (Figure 1, b). The formation of a 1:1 complex is coincident with maximal Sirt7 activation towards H3K18Deca peptide, which rapidly declines with the formation of the second low affinity 2:1 complex dominating at a high 8:1 Sirt7/tRNA ratio. Similar to the observation with 146bp DNA, further increasing Sirt7 concentration resulted in aggregation and a loss of distinct bands revealed by the EMSA analysis.

The addition of the Sirt7 to nucleosomes initially results in the simultaneous formation of predominantly 1:1 and 2:1 Sirt7/nucleosome stoichiometric complexes (Figure 1 c). This behavior is consistent with specific Sirt7 interactions with two symmetrical acidic patch^24,25^ regions on the nucleosomal surface, which was previously observed with the highly homologous Sirt6.^26^ To confirm that the nucleosome acidic patch is important for Sirt7 binding, we synthesized a FAM-labeled LANA peptide probe^27^ known to specifically bind to the acidic patch regions of nucleosomes (Figure 1, d, e).^28^ Competitive displacement of LANA-FAM peptide from the pre-formed LANA- nucleosomal complex by Sirt7 was monitored by the increase of fluorescence anisotropy (Figure 1, f). It supports a mechanism of binding where Sirt7 competes with the labeled LANA probe for the same negatively charged acidic patch regions (Figure 1, a). Unlike Sirt6 which only forms 1:1 and 2:1 stoichiometric complexes,^26^ an additional distinct 3:1 Sirt7/nucleosome complex was detected by EMSA at a higher Sirt7/nucleosome ratio suggesting another but weaker interaction. Interestingly, the addition of Sirt7 to nucleosomes gradually increases the long-chain H3K18Deca deacylation activity up to 8:1 Sirt7/nucleosome ratio. This activation response is similar to the observation with free 146 bp DNA (Figure 1, a) that indicates the disruption of nucleosome core particles by Sirt7 with concomitant formation of soluble heterogeneous aggregates revealed by corresponding EMSA. Further reduction of total activity occurs at a higher 10:1 Sirt7/nucleosome ratio that is coincident with additional aggregation.

The conducted EMSA analysis of Sirt7 interacting with DNA, tRNA, and nucleosomes provided valuable novel information regarding the behavior of complexes, the size distribution, and catalytic activity. Overall, Sirt7 uniquely interacts with DNA, tRNA, or nucleosomes resulting in activation of the long-chain deacylation activity. In all cases, a specific 1:1 stoichiometric complexation was sufficient for maximal catalytic activation of each Sirt7 equivalent. Because interaction with excessive Sirt7 equivalents caused aggregation and reduction of catalytic activity, all subsequent studies were conducted at a 1:1 stoichiometric ratio to allow a more accurate molecular assessment of Sirt7 activity in the presence of various modulators.

### Minimal nucleic acid interaction footprint for Sirt7 activation

The EMSA analysis revealed that 146 bp-long DNA at a 1:1 Sirt7/DNA ratio (Figure 1, a) forms multiple nucleoprotein species, while tRNA at the same stoichiometric ratio only forms a single 1:1 stoichiometric complex (Figure 1, b). Long heterogeneous DNA fragments were previously reported to be inferior activators of Sirt7 in comparison with tRNA.^29^ To properly evaluate the relative activating potential of DNA, tRNA, and nucleosomes and to avoid potential complications from the formation of heterogeneous poly-substituted Sirt7-DNA species, we tested the activating potential of short double-stranded oligo- DNAs (Supplementary Table S1) on purified recombinant apoSirt7. Considering that each molecule of Sirt7 interacts with 18-20 bp long regions of the 146 bp DNA fragment (Figure1, a), we conducted Michaelis-Menten kinetic analysis of Sirt7 in the presence of short synthetic double-stranded oligo-DNAs (Figure 2, a).

**Figure 2.**
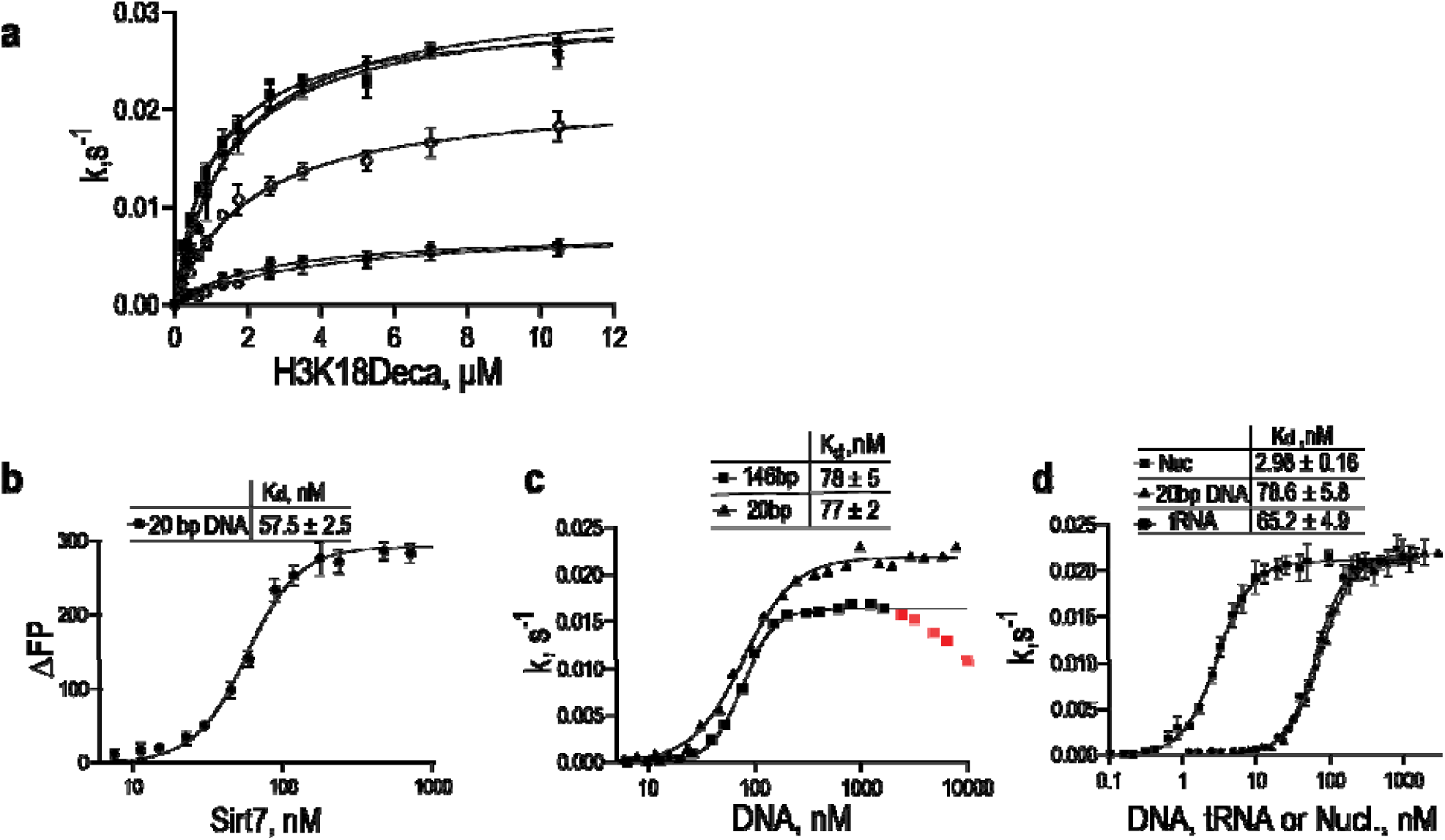
Sirt7 interaction and activation by short double-stranded oligo-DNA, tRNA, and nucleosomes. a. Michaelis-Menten profiles of Sirt7 H3K18Deca deacylation in the presence of various activator DNAs and corresponding kinetic parameters (Table1). b. Fluorescence polarization (FP) analysis of Sirt7 interaction with 20bp double-stranded FAM-labeled DNA. c. Kinetic titration of long 146bp DNA and short 20bp DNA into Sirt7 in the presence of saturating substrates. d. Titration of 20bp double-stranded DNA, tRNA, and nucleosomes into Sirt7 in the presence of saturating substrates.

This kinetic analysis revealed that Sirt7 achieves maximal catalytic efficiency (k_cat_/K_M_, Table 1) in complex with 15-20 bp double-stranded oligo-DNAs. Thus, considering the average length of 0.34 nm/bp (3.4 Å) the size of the minimal footprint of the double-stranded oligo-DNA required for Sirt7 activation was calculated to be 5-7 nm (50-70 Å). This value matches previously reported linear dimensions of average folded tRNA molecules.^30^ The fact that the linear dimensions of compactly folded 75-90 bp long tRNA molecule match the size of the minimal Sirt7-activating DNA footprint explains the predominant formation of the single 1:1 stoichiometric Sirt7-tRNA complex observed by EMSA (Figure 1, b). Consistently, a sterically unfavorable mutually interfering interaction of two Sirt7 equivalents with one folded tRNA molecule observed by EMSA is weaker and results in reduced activation compared to a more favorable and specific 1:1 stoichiometric complex (Figure 1, b). The evidence also suggests that the 3D arrangement and nature of the nucleic acids may not significantly impact Sirt7 binding affinity or activation as long as the linear dimensions of the negatively-charged polyanionic species match the minimum size requirement.

**Table 1.**
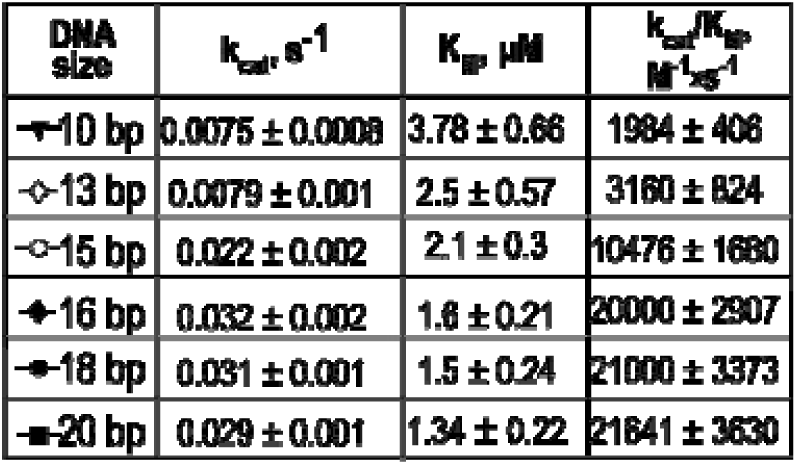
Sirt7 activation by short dsDNA

### Binding affinity and activating potential of various Sirt7 modulators

To further evaluate the effect of size, folding, and biological nature of various modulators we tested short and long DNAs, tRNA, and nucleosomes for the ability to bind and activate apoSirt7. First, we conducted a fluorescence polarization (FP) binding experiment using 20 bp FAM-labeled DNA probe. We found that this short oligo-DNA effectively interacts with Sirt7 with K_d_ = 57.2 nM (Figure 2, b). To further validate the binding affinity and at the same time to evaluate the limiting activating potential, we titrated Sirt7 with long (146bp) or short (20bp) double-stranded DNAs in the presence of saturating NAD^+^ and long-chain H3K18Deca acylated peptide (Figure 2, c). The kinetic titration of DNAs into Sirt7 revealed that the enzyme equally well binds long or short DNA, and both interactions result in similar activation (Figure 2, c). However, the enzymatic activity was clearly inhibited by high concentrations of long, but not the minimal 20bp DNA (Figure 2, c). The inhibition of Sirt7 activity by aggregation was not accounted for in previous *in vitro* studies,^19^ likely resulting in a significant underestimation of Sirt7-DNA complex activity and ultimately in the conclusion that tRNA provided superior activating potential.^22^Thus, unlike previous reports that utilized a large excess of long sonicated heterogeneous salmon sperm DNA for Sirt7 activation^19^ we chose to use only short synthetic HPLC-purified 20 bp double-stranded oligo-DNA in our Sirt7 comparative enzymatic analysis.

To directly compare the binding affinity and activating potential of all modulators, 20 bp DNA, tRNA, and nucleosomes were individually titrated into Sirt7 in the presence of saturating NAD^+^ and long-chain acylated H3K18Deca peptide substrate (Figure 2, d). Analysis of the resulting binding curves revealed that despite previous report^29^ the tRNA has no activating or binding advantage over short linear 20bp DNA (Figure 2, c).

Interestingly, titration of nucleosomes into Sirt7 under the same conditions revealed a 30-fold stronger interaction (lower K_d_) with a similar degree of activation towards the long-chain acylated substrate compared to all nucleic acids. The much tighter affinity indicates a strong preference of Sirt7 to bind nucleosomes over DNA or tRNA. To further investigate the details of Sirt7 interaction with DNA, tRNA, and nucleosomes we developed a direct nucleosome binding assay based on the NanoLuc luciferase complementation principle.

### Design and application of NanoLuc complementation assay

The discovery of nucleosomes as a low nanomolar high affinity Sirt7 binding partner prompted us to develop a novel specialized tool to further investigate the mechanistic aspects of the Sirt7-nucleosome interaction as well as the enzyme preference to interact with various modulators.

In this work, we report a successful development and application of a highly sensitive *in vitro* NanoLuc split luciferases binding assay^31^ for quantitative direct measurement of sirtuin interaction with nucleosomes (Figure 3, a). This direct nucleosome binding assay complements the above kinetic approach (Figure 2, d), but features much higher sensitivity and does not depend on sirtuin activity response, thus allowing measurements of Sirt7 affinity for unmodified as well as acetylated nucleosomes.

**Figure 3.**
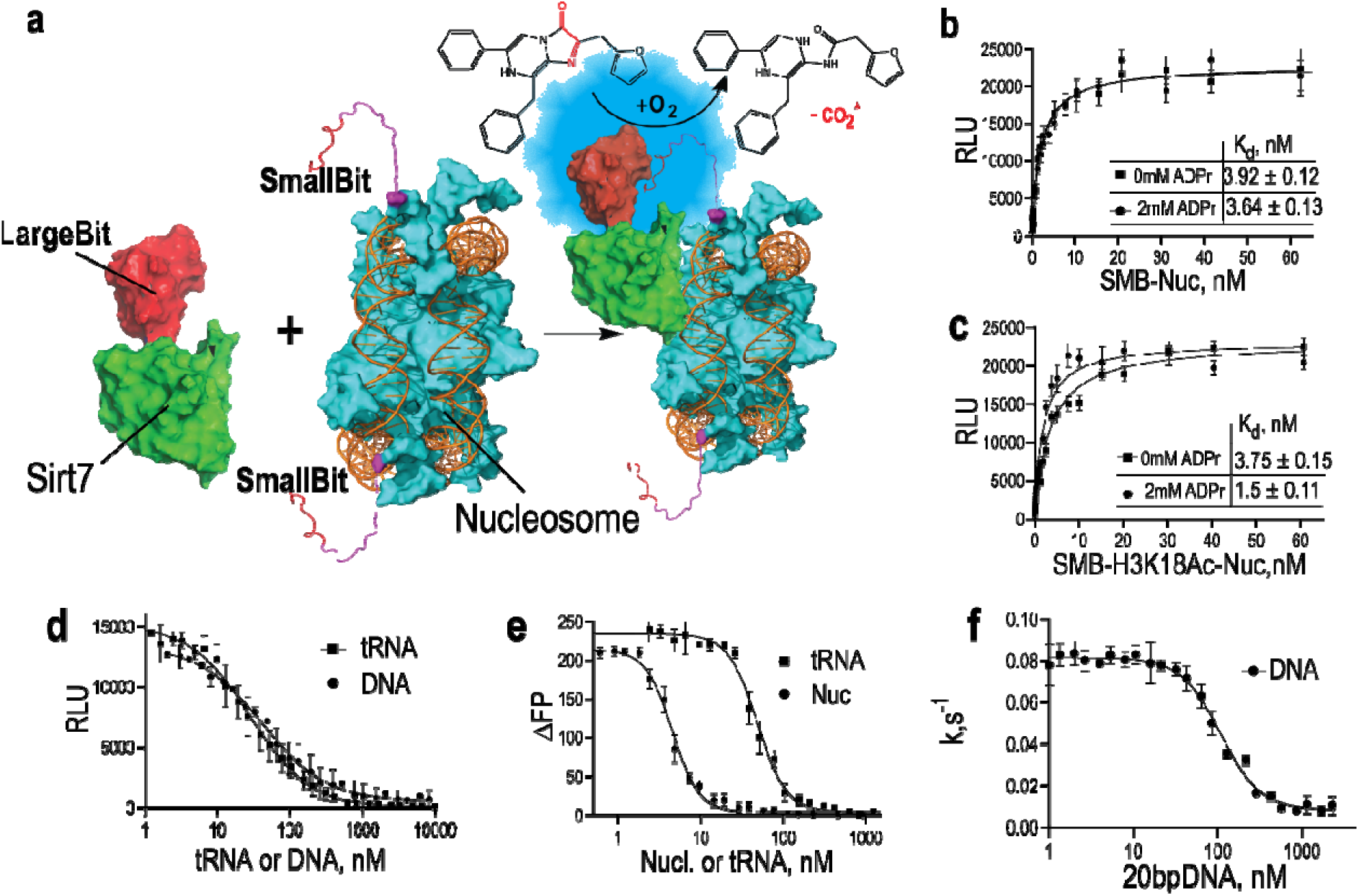
Development and application of NanoLuc nucleosome binding assay. a. The principle of the novel in vitro NanoLuc complementation binding assay designed to study Sirt7-nucleosome interaction. b, c. Effect of ADP-ribose on Sirt7 binding affinity to unmodified and H3K18-acetylated nucleosomes, respectively. d. NanoLuc luminescence decay associated with competitive displacement of Sirt7- CLB from SMB-Nucs by 20bp DNA and tRNA. e. Fluorescence anisotropy reduction associated with competitive displacement of 20bp FAM-DNA from Sirt7 by nucleosomes and tRNA. f. Inhibition of Sirt7 deacetylation activity by DNA. Reduction of Sirt7 deacetylation activity associated with competitive displacement of nucleosomes by 20bp DNA in the presence of H3K18Ac peptide.

Because the preparation of nucleosomes from recombinant components (histones and DNA) requires a chaotropic denaturation, we chose to attach the unstructured small-bit (SMB) peptide tag to the N-terminus of histone H2B. This site-specific nucleosome epitope tagging approach does not affect the tertiary structure and is well tolerated by chromatin binding proteins. The complementary large bit component was fused to the C-terminus of Sirt7 (Sirt7-CLB) and purified under native conditions. The principal nature of the assay (Figure 3, a) relies on the primary Sirt7-nucleosome interaction that increases the frequency of NanoLuc luciferase complementation events thus resulting in a drastic increase in luminescence signal. The isolated large bit domain alone very weakly (K_d_>5 µM) interacts with the SMB-Nucs (Supplementary figure S3), which does not affect the targeted interaction of tight nucleosome binding proteins such as Sirt7.

The unmodified and H3K18 acetylated small bit nucleosomes (SMB-Nucs) were titrated into Sirt7-CLB to determine K_d_ values for 1:1 stoichiometric complex formation (Figure 3, b, c). Because ADP-ribose (ADPr) binding was previously shown to enhance Sirt6 affinity for its acetylated peptide substrate,^32^ we chose to include this NAD^+^ analog in our study. In the absence of ADPr, unmodified and acetylated nucleosomes interacted with the same affinity. The measured K_d_ values (Figure 3, b, c) of 3.92 nM and 3.75 nM agree well with the binding K_d_ constant of 2.92 nM obtained above by the kinetic titration (Figure 2,d). However, the 2.5-fold reduction in K_d_ value was observed when ADPr was included with H3K18-acetylated, but not unmodified nucleosomes. This data suggests that the interaction of ADPr with the Sirt7-nucleosome complex precedes and promotes the additional interactions of the H3K18-acetylated tail with the active site.

In addition, the NanoLuc assay was used to investigate whether nucleic acid and nucleosomes directly compete or interact with orthogonal binding sites on the surface of Sirt7. As Sirt7 interacts with nucleosomes with 40-fold higher affinity compared to free nucleic acids, it is mechanistically important to evaluate the mutual competition between these poly-anionic modulators. In the nucleus, the ability of nucleosome-bound Sirt7 to additionally interact with free nucleic acids would allow for DNA and RNA to further modulate its enzymatic activity and biological functions. Thus, we conducted a series of direct competition experiments by monitoring equilibrium dissociation of Sirt7- nucleosome (NanoLuc) or Sirt7-DNA(FAM) complexes (Figure 3, d, e). The data indicate that DNA, tRNA, and nucleosomes are mutually exclusive and directly compete for an overlapping binding site on the surface of Sirt7. Consistently with the affinity difference, the stoichiometric addition of nucleosomes to Sirt7-DNA complex completely displaces the nucleic acid (Figure 3, e) while 1000-fold excess of nucleic acids (DNA or tRNA) was required to break the Sirt7-nucleosome complex (Figure 3, d). The large Sirt7 preference to bind nucleosomes and its inability to interact with nucleosomes and free nucleic acids at the same time emphasizes the higher biological importance of nucleosomes. At the same time, it implies a complex multidentate Sirt7-nucleosome binding mechanism that involves both nucleosomal DNA and the core histone octamer. Indeed, the direct competition between nucleic acids and nucleosomes as well as competitive displacement of the LANA fluorescent probe from the acidic patch by Sirt7 (Figure 1, d-f) indicate that the enzyme simultaneously interacts with nucleosomal core DNA and the histone octamer. From the standpoint of *in vivo* function, the enormous excess of free nucleic acids required to displace nucleosome-bound Sirt7 suggests that in the presence of chromatin the cellular Sirt7 even in the nucleolar RNA-rich compartment^5^ will tend to interact with nucleosomes.

Because Sirt7-DNA and Sirt7-RNA complexes previously displayed weak deacetylase activity^19,29^ and to evaluate the functional effect of the nucleosome and nucleic acid competition on enzymatic activity, we measured Sirt7 kinetic response by titrating short 20 bp DNA into pre-formed Sirt7-nucleosome complex in the presence of H3K18Ac peptide substrate and saturating NAD^+^. Titration of oligo-DNA to nucleosome pre-activated Sirt7 resulted in dose-dependent inhibition of deacetylation activity (Figure 3, f). Thus, the interaction of Sirt7 with nucleosomes appears to be particularly effective for activation of Sirt7 deacetylation activity, while for the long-chain deacylation activity both nucleosomes and nucleic acids are equally effective (Figure 2, d).

### Optimization of sirtuin kinetic deacetylation assay

To further investigate the role of various modulators on Sirt7, we improved the existing UV-HPLC-based peptide assay^33^ that remains one of the most reliable, direct, and commonly used methods for the evaluation of *in vitro* sirtuin activity. The major downsides of the original UV-HPLC methodology are long separation gradients, time-consuming data processing, and high UV-interfering background caused by numerous large and small molecular weight assay components such as NAD, NAM, DMSO, DNA, RNA, Sirt7, minor peptide impurities, and histones. Because Sirt7 displayed the strongest peptide K_M_ values in the sirtuin family the high UV background associated with the presence of interfering components severely complicated analysis with the conventional tryptophan labeled peptides that allows detection only in the UV spectral range. To overcome the issues and improve the high-throughput capability of the assay, we synthesized our H3K18-acylated Sirt7 substrates using an alternative 1,4-dinitrofluorobenzezene (DNB, Sanger reagent) labeling strategy to substitute for conventionally used tryptophan (Figure 4, a).

**Figure 4.**
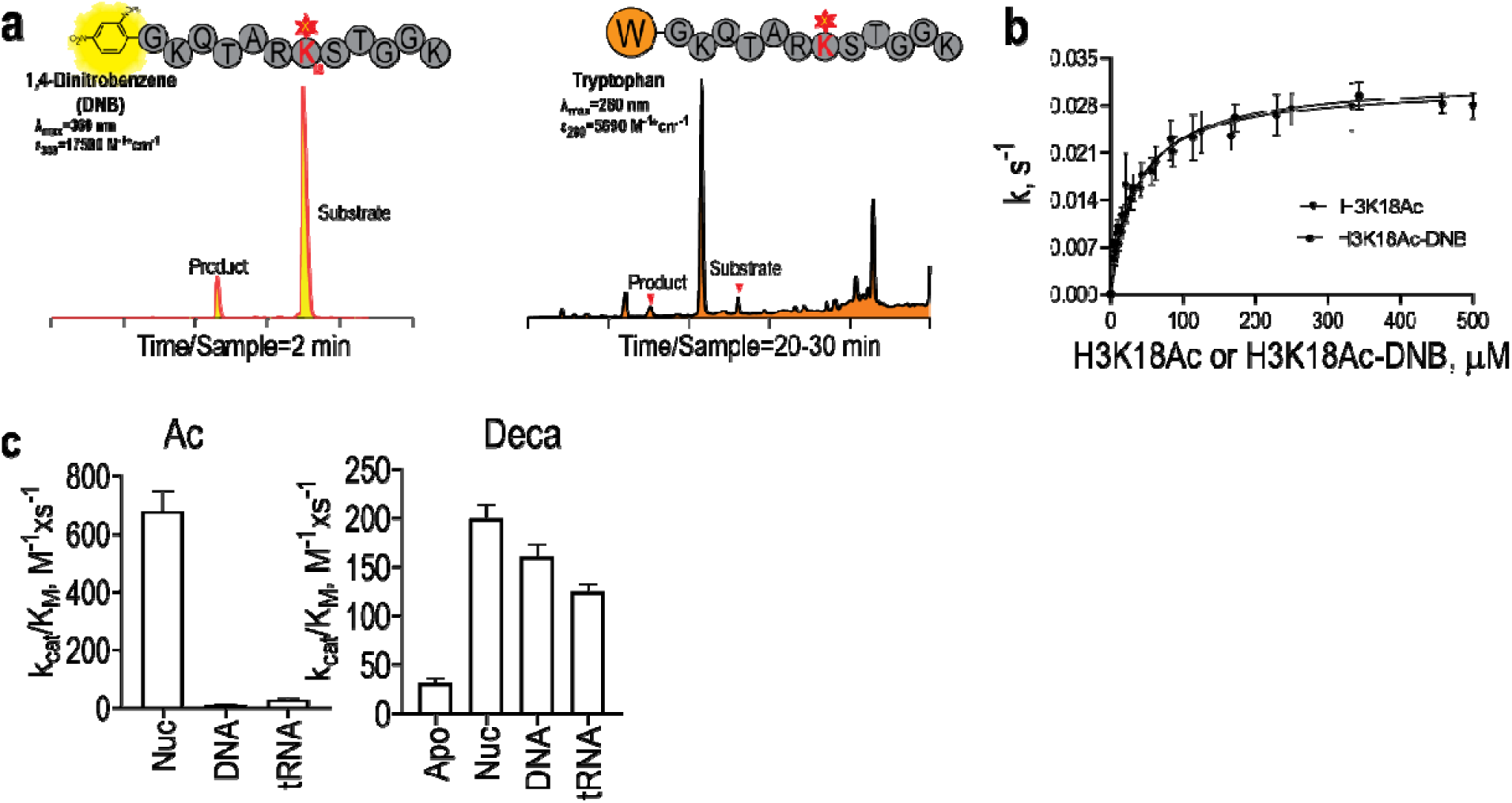
Design and validation of novel HPLC-based sirtuin deacetylation assay. a. Sirtuin deacetylation assay optimization concept that improves its sensitivity and high-throughput capability. HPLC separation profiles of Sirt7-deacetylated 1.4- dinitrobenzene (DNB) and tryptophan (W) labeled peptides. b. Validation of DNB- labeled Sirt7 substrate demonstrates that its kinetic behavior is unaffected by the peptide labeling strategy. c. Effect of DNA, tRNA, and nucleosomes on Sirt7 NAD^+^- catalytic efficiencies in the presence of saturating H3K18Ac or long-chained H3K18Deca substrates. The Sirt7 kinetic parameters are summarized in the table, and corresponding Michaelis-Menten profiles are shown in Supplementary figure S4.

The new labeling approach did not affect the kinetic properties of Sirt7 (Figure 4, b), but allowed monitoring reaction at 400-450 nM and thus effectively eliminated the original UV interference from other molecules in the assay. Due to reduced sample runtime and greatly simplified data processing, the optimized HPLC assay can be more effectively utilized for high-throughput analysis of sirtuin enzymatic activity.

### Sirt7 kinetic analysis with NAD^+^ as a variable substrate

To investigate the enzymatic aspects of Sirt7 catalysis that are affected by DNA, tRNA, and nucleosomes, the optimized HPLC assay was used to determine kinetic parameters in the presence of saturating H3K18Ac or H3K18Deca peptides and variable NAD^+^(Supplementary figure S4). Compared to apoSirt7 with H3K18Deca peptide, the complexation with nucleic acids or nucleosomes significantly improved the enzymatic efficiency by reducing K_M_^NAD^ values without any significant effect on k_cat_ (Figure 4, c, Table 2). However, with the H3K18Ac substrate, the activating effect of nucleosomes was significantly stronger compared to DNA and tRNA (Figure 4, c). K_M_^NAD^ values for nucleosome activated Sirt7 were the same for both H3K18Ac and H3K18Deca peptide substrates. However, the nucleic acid activation was strongly dependent on the length of the H3K18 peptide acylating group. With H3K18Ac substrate peptide, K_M_^NAD^ values for Sirt7 activated by DNA and tRNA were higher by 50- and 20-fold, respectively, compared to nucleosomes that resulted in a significant reduction of Sirt7 catalytic efficiency in the presence of nucleic acids (Figure 4, c). From the standpoint of *in vivo* function, the elevated K_M_^NAD+^ values of 0.48 mM and 1.27 mM for Sirt7-tRNA and Sirt7-DNA complexes, respectively in comparison to 0.1 mM K_M_^NAD+^ for Sirt7 bound to nucleosomes suggests the limited functionality of Sirt7-nucleic acid complexes toward acetylated targets. Considering that physiological NAD^+^ level ranges from 0.1-0.5 mM^34^ the deacetylation efficiency of Sirt7- DNA and Sirt7-tRNA complexes would be naturally limited by the availability of the co-substrate in the cell. However, outside of the nuclear envelope and in the absence of competing nucleosomes, free nucleic acids may still have a significant biological effect on Sirt7 with cytoplasmic long- chain acylated targets that lower K_M_^NAD+^ values to the physiologically relevant range.

**Table 2.**
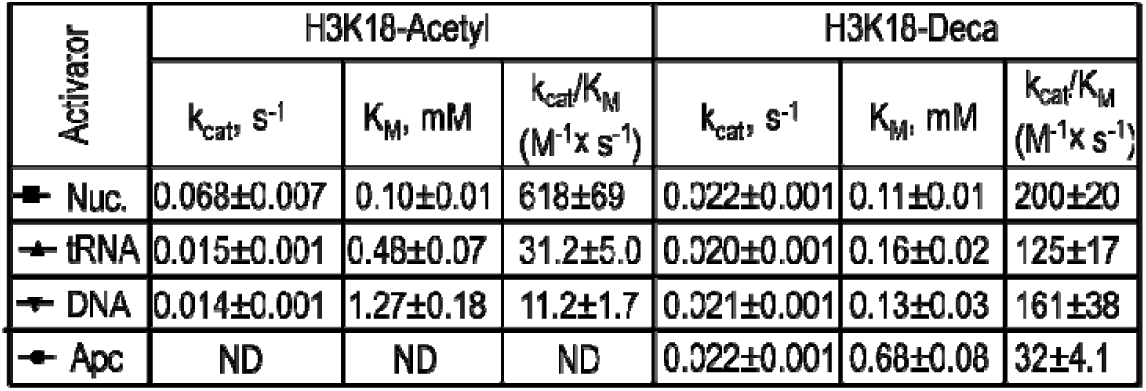
NAD Kinetic Parameters.

### Sirt7 kinetic analysis with various H3K18 acylated peptides as variable substrates

Kinetic analysis with variable NAD^+^ substrate strongly suggested that Sirt7 activation by nucleosome or nucleic acid modulators is dependent on the nature of the peptide substrate. From previous work,^33^ different members of the sirtuin family displayed distinct preferences for short- or long-chain acylated peptides. While the chemical nature of the lysine acylating group was known to control the level of sirtuin catalytic efficiency and the scope of enzymatic substrates,^33^ these target preferences had never been studied in combination with additional modulators such as nucleic acids or nucleosomes that frequently co-localize with sirtuins in the cellular environment. Complexation with nucleic acids or nucleosomes represents an additional way of Sirt7 regulation that may contribute to or work independently of the activation by hydrophobic substrate acylating groups. To explore this, a systematic experimental analysis was conducted in the presence of nucleic acids and nucleosomes using a broad spectrum of short- and long-chain hydrophobic H3K18-modified substrates (Figure 5 a). First, the effect of substrate acylation length (hydrophobicity) on uncomplexed SIRT7 (apoSirt7) was determined. In contrast to previous reports^19,29^ that apoSirt7 was completely inactive, we determined a very low, but readily measurable H3K18 deacetylation activity (Figure 5, a). Substituting H3K18Ac with a long-chain hydrophobic H3K18Deca substrate resulted in a significant rate enhancement as evidenced by a 10-fold increase of k_cat_ and a 40-fold improvement of K_M_ values, which overall translated into a 585-fold increase of the apoSirt7 catalytic efficiency (Figure 5, a).

**Figure 5.**
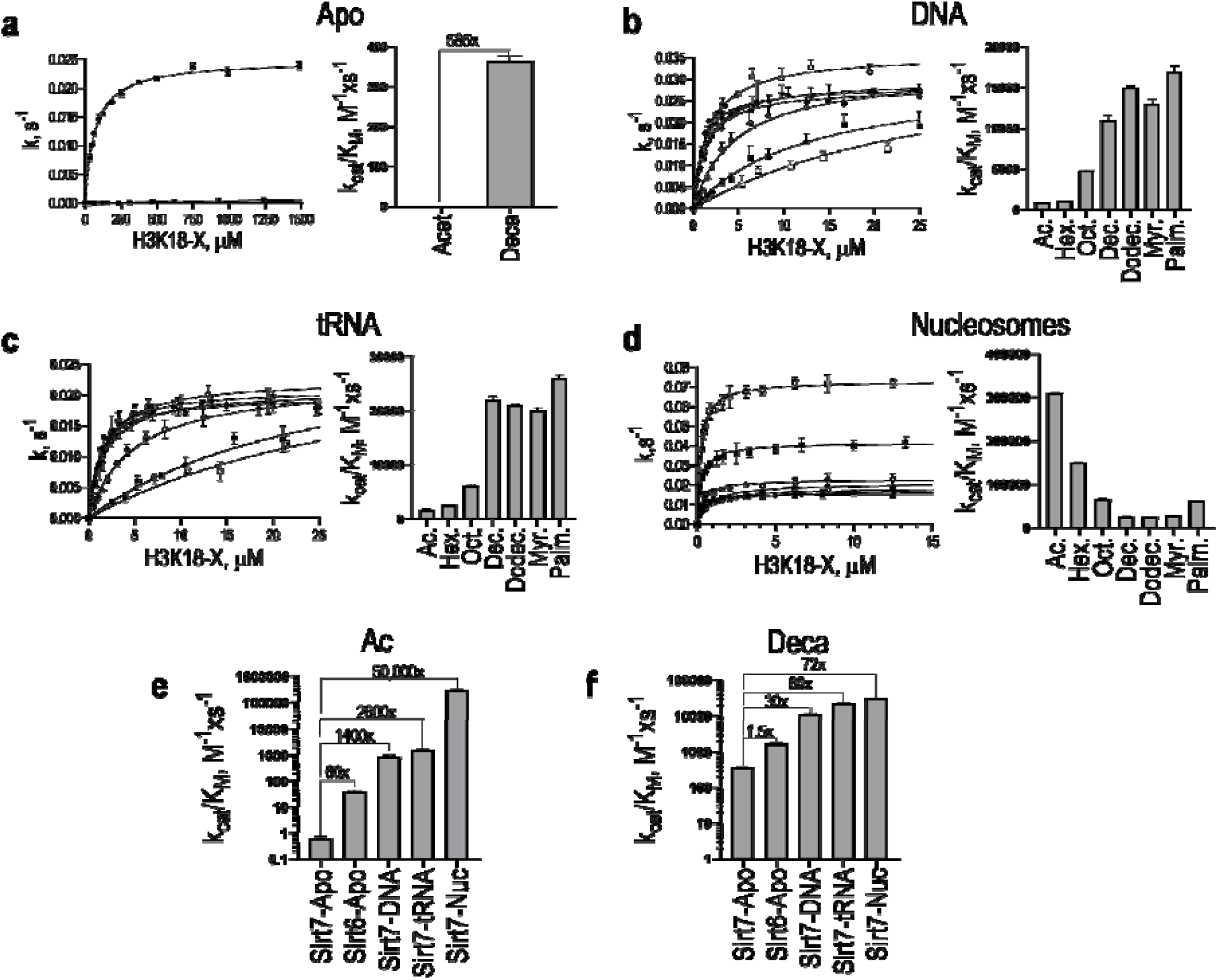
Effect of modulators on Sirt7 kinetics with various H3K18 acylated substrates. Michaelis-Menten profiles and bar plots of corresponding catalytic efficiencies in the absence of modulators (a), and in the presence of DNA (b), tRNA (c), or nucleosomes (d). The values of catalytic efficiencies (Table 3) of apoSirt7, apoSirt6, Sirt7-DNA, Sirt7-tRNA, and Sirt7-nucleosome complexes towards H3K18Ac (e) and H3K18Deca (f) substrates presented as bar plot on the vertical logarithmic

This hydrophobic substrate-mediated rate enhancement was previously reported for highly homologous apoSirt6^33^ that relatively to apoSirt7 was 60 fold more efficient with H3K18Ac (Figure 5, e) and equally efficient towards H3K18Deca substrate (Figure 5, f). Complexation with nucleic acids improves the Sirt7 deacetylation efficiency 1400 and 2600 times over nearly inactive apoSirt7 while the respective long-chain deacylation efficiency only increases 30- and 60-fold for DNA and tRNA, respectively (Figure 5, e, f). However, the resulting Sirt7-DNA or Sirt7-tRNA complexes still display very poor enzymatic efficiencies towards the H3K18Ac substrate as evidenced by the extremely high K_M_^Ac^ and flat peptide saturation curves (Figure 5, b, c). By comparison, Sirt7 interaction with nucleosomes results in 50000-fold activation towards H3K18Ac and 70-fold activation towards H3K18Deca deacylation (Figure 5, e, f). Steep hyperbolic Michaelis-Menten profiles (Figure 5, d) are observed in the presence of nucleosomes for the entire range of H3K18 short- and long-chain acylated peptides. Although both nucleic acids and nucleosomes equally well activate long-chain deacylation activity (Figure 5, f), the DNA and tRNA activation effect rapidly decreases as the length of the H3K18 acylating group decreases (Figure 5, b, c, d). In contrast, the catalytic efficiency of Sirt7 activated by nucleosome further improves with shorter acyl modification (Figure 5, d). Collectively, these results strongly suggest that nucleosomes are ultimate universal activators of Sirt7, capable of activating both short- and long-chain deacylase activity, unlike complexation with DNA or tRNA that provides only partial Sirt7 activation that is strongly dependent on the peptide substrate properties. When properly activated by nucleosomes, all Sirt7 kinetic parameters significantly improve suggesting that consistently with numerous biological reports the scope of enzymatic activity is not limited to hydrophobic long-chain acylated substrates and Sirt7 is a highly functional deacetylase in the cell.

### *Ex vivo* chromatin deacetylation by Sirt7

To test the Sirt7 deacetylation efficiency towards cellular substrates we conducted a controlled activity assay using purified native chromatin (oligo-nucleosomes) isolated from mammalian cells. We prepared native oligo-nucleosomes using our published methodology.^35^ The protocol allows isolation of native chromatin fragments chromatographically separated from contaminating enzymatic and non-enzymatic impurities while preserving combinatorial PTMs.^35^ Sirt7 activity was investigated using substrate oligo-nucleosomes purified from histone deacetylase (HDAC) inhibitor-treated PC3 cancer cells (Figure 6, a). Recombinant Sirt6 was chosen as a specific positive control for its known high activity and selectivity towards H3K9Ac in the context of the nucleosomes.^36^ Sirt2 on the other hand was utilized as a non-specific positive control to evaluate the global sirtuin accessibility to all acetylated sites.^37^ Deacetylated chromatin was analyzed using both LC-MS/MS (Figure 6, b) and western dot-blot (Figure 6, c) with anti-K9Ac and anti-K18Ac antibodies. Sirt7 displayed a high degree of selectivity by specifically deacetylating H3K18Ac and H3K27Ac chromatin marks.

**Figure 6.**
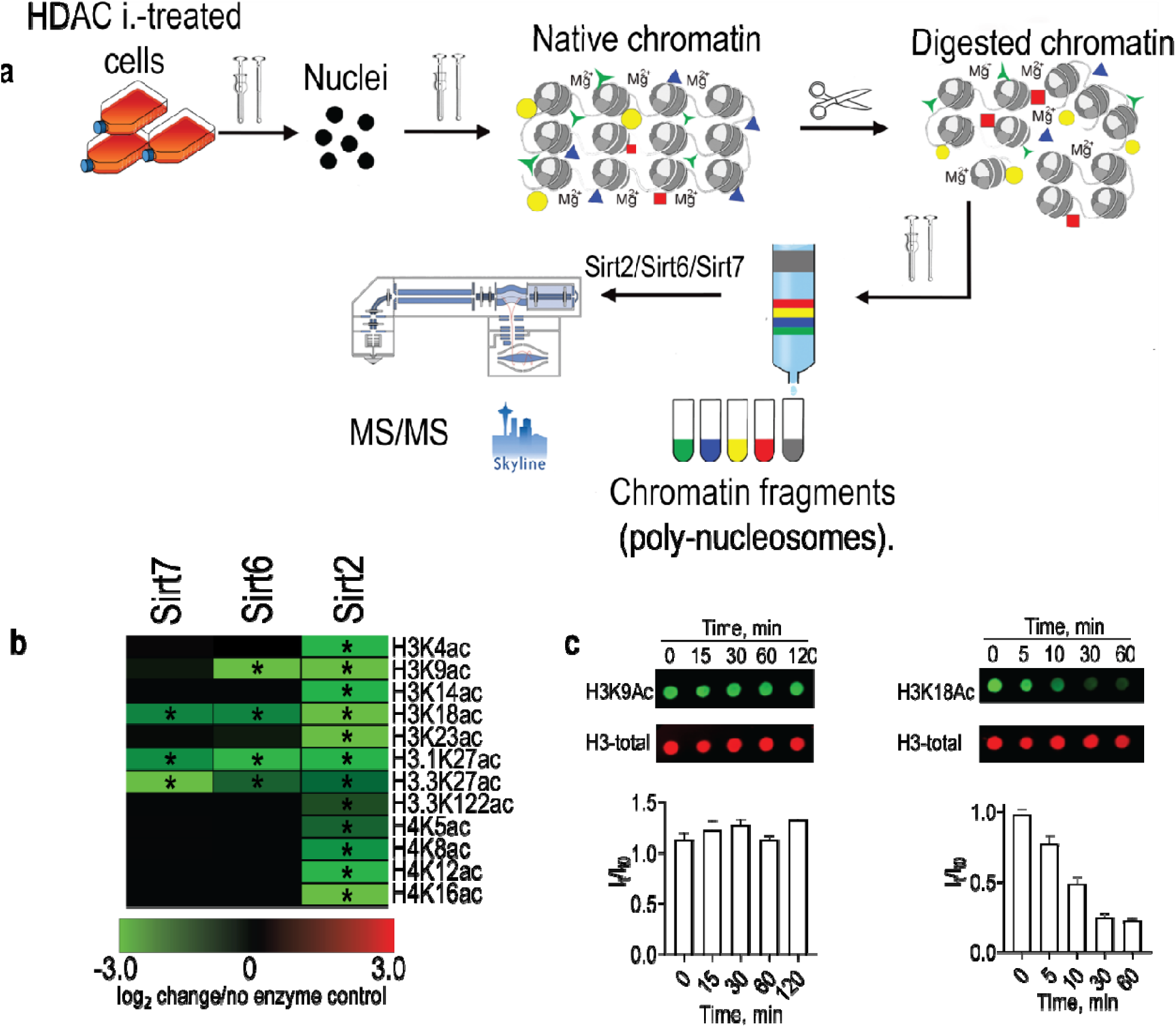
Design and analysis of Sirt7 ex vivo chromatin deacetylation specificity. Experimental design (a) and (b) LC/MS analysis results of chromatin deacetylation by Sirt2, Sirt6, and Sirt7 presented as a heat-map relative to untreated control. c. The time course of Sirt7 chromatin deacetylation by western dot-blot with anti-H3K9Ac and anti-H3K18Ac antibodies.

Sirt6 displayed less specificity and in addition to the generally accepted H3K9Ac deacetylation was highly efficient towards H3K18Ac and H3K27Ac modifications. Sirt2 removed all acetylation marks, consistently with prior knowledge as a more promiscuous NAD^+^-dependent deacetylase.^37^ These results demonstrate that consistently with previous biological studies^21^ the nucleosome-activated Sirt7 is a highly site-specific and catalytically efficient chromatin deacetylase.

### Intramolecular Sirt7 catalytic deacetylation efficiency

To further demonstrate that Sirt7 can deacetylate the directly associated substrate nucleosome, and to quantitatively evaluate the catalytic efficiency of this intramolecular deacetylation process we designed a novel antibody-free single-turnover kinetic experiment (Figure 7, To specifically determine the rates of Sirt7 intramolecular nucleosome deacetylation towards pre-validated H3K18Ac or H3K27Ac chromatin sites, we monitored the formation time course of acetyl-ADP-ribose (AcADPr) and nicotinamide (NAM) products using LC/MS. Recombinant site-specifically acetylated nucleosomes were assembled with minimal 146bp Widom 601 DNA to additionally demonstrate that in contrast to previously reported hypothesis^19^ the nucleosome core particle itself, and not the linking DNA activates Sirt7 deacetylation activity. Pre-saturating acetylated substrate nucleosomes with Sirt7 prior to addition of NAD^+^ allows monitoring single turnover intramolecular deacetylation process by measuring NAM and AcADPr formation that report the rates *O*-alkyl-amidate formation and breakdown, respectively (Figure 7, a). The first order intramolecular deacetylation rates of Sirt7-catalyzed deacetylation of H3K18 and H3K27 acetylated nucleosomes (Figure 7, b) are k^NAM^=0.024 and 0.033 s^-1^, and k^AcADPr^=0.023 and 0.039 s^-1^, respectively. The rates are very similar to the steady-state k_cat_ values determined for intermolecular deacetylation of free H3K18Ac peptide by Sirt7, pre-activated with unmodified nucleosomes (Figure 5, Table 3). However, to determine the Sirt7 catalytic efficiency (k_cat_/K_M_) for the multiple turnover intramolecular deacetylation of acetylated nucleosomes, the dissociation of Sirt7 from unmodified deacetylated nucleosomes must be measured as the product release contributes to both k_cat_ and K_M_ kinetic parameters (Figure 7,d). To measure the rates of Sirt7 dissociation from the substrate and product nucleosomes (k_off_^AcNu^ and k_off_^Nu^, Figure 7, d) we used the NanoLuc dissociation assay. In the presence of 1000- fold excess of untagged nucleosomes, the dissociation rate of Sirt7-CLB from the SMB- containing unmodified or K18 acetylated nucleosomes was measured by monitoring the first order luminescence signal decay (Figure 7, c). Using previously measured equilibrium binding constants (Figure 3, b, c) the remaining individual rate constants of the intramolecular steady-state nucleosome deacetylation process can be easily calculated (Figure 7, d). The measured Sirt7-nucleosome dissociation rate is 10 times slower compared to the rates of all chemical steps k_chem_ (k^NAM^ or k^AcADPr^). Thus, the product (unmodified nucleosomes) dissociation rate becomes limiting and determines the k_cat_ value for the Sirt7-catalyzed multiple turnover intramolecular nucleosome deacetylation process (k_cat_)^AcNu^= (k_off_)^Nu^=0.0042 s^-1^). Despite 17 fold lower (k_cat_)^AcNu^=0.0042 s^-1^, compared to the steady-state intermolecular H3K18Ac peptide deacetylation (k_cat_=0.073 s^-1^, Table 3) the (K_M_)^AcNu^=(k_off_^AcNu^ +k_cat_)/(k ^AcNu^) =3.2×10^−9^ M for intramolecular deacetylation process is 72-fold better resulting in further improvement of Sirt7 catalytic efficiency (k_cat_/K_M_)^AcNu^=0.13×10^7^ (M^-1^*s^-1^). It is moderately (4 times) higher than the corresponding intermolecular H3K18Ac peptide deacetylation (k_cat_/K_M_=0.317×10^6^, Table 3) by nucleosome-activated Sirt7, but greatly exceeds the corresponding intermolecular catalytic efficiencies of Sirt7 activated by DNA and tRNA (k_cat_/K_M_=0.09 ×10^4^ M^-1^*s^-1^ and k_cat_/K_M_=0.16×10^4^ M^-1^*s^-1^, respectively, Table 3). Altogether, these results indicate that Sirt7 is a highly specific and efficient intra- and intermolecular deacetylase in complex with nucleosomes, while interaction with DNA or tRNA is much weaker and insufficient for activation of Sirt7 deacetylation activity.

**Table 3.** Peptide kinetic parameters (saturating NAD)

| H3K18-X | DNA |  |  | tRNA |  |  | Nucleosomes |  |  | Apo |  |  |
| --- | --- | --- | --- | --- | --- | --- | --- | --- | --- | --- | --- | --- |
| | $k_{cat}$<br>($s^{-1}$) | $K_M$<br>($\mu M$) | $(k_{cat}/K_M) \times 10^3$<br>($M^{-1} s^{-1}$) | $k_{cat}$<br>($s^{-1}$) | $K_M$<br>($\mu M$) | $(k_{cat}/K_M) \times 10^3$<br>($M^{-1} s^{-1}$) | $k_{cat}$<br>($s^{-1}$) | $K_M$<br>($\mu M$) | $(k_{cat}/K_M) \times 10^3$<br>($M^{-1} s^{-1}$) | $k_{cat}$<br>($s^{-1}$) | $K_M$<br>($\mu M$) | $(k_{cat}/K_M) \times 10^3$<br>($M^{-1} s^{-1}$) |
| Acetyl | $0.031 \pm 0.001$ | $34.5 \pm 4.9$ | $0.9 \pm 0.13$ | $0.030 \pm 0.0017$ | $18.36 \pm 1.96$ | $1.69 \pm 0.2$ | $0.073 \pm 0.002$ | $0.23 \pm 0.04$ | $317 \pm 56$ | $0.0018 \pm 0.0003$ | $2864 \pm 508$ | $0.00063 \pm 0.00015$ |
| Hexanoyl | $0.026 \pm 0.002$ | $25.5 \pm 4.1$ | $1.1 \pm 0.19$ | $0.027 \pm 0.0025$ | $10.76 \pm 1.97$ | $2.56 \pm 0.51$ | $0.041 \pm 0.001$ | $0.27 \pm 0.04$ | $152 \pm 23$ | ND | ND | ND |
| Octanoyl | $0.022 \pm 0.001$ | $4.54 \pm 0.61$ | $4.8 \pm 0.9$ | $0.025 \pm 0.0016$ | $4.76 \pm 0.73$ | $5.1 \pm 1.9$ | $0.022 \pm 0.001$ | $0.34 \pm 0.11$ | $65 \pm 21$ | ND | ND | ND |
| Decanoyl | $0.021 \pm 0.001$ | $1.67 \pm 0.26$ | $11.2 \pm 1.8$ | $0.020 \pm 0.0017$ | $1.32 \pm 0.28$ | $21.9 \pm 4.8$ | $0.021 \pm 0.001$ | $0.8 \pm 0.14$ | $26.2 \pm 4.7$ | $0.025 \pm 0.001$ | $68.8 \pm 5.4$ | $0.36 \pm 0.03$ |
| Dodecanoyl | $0.023 \pm 0.001$ | $1.34 \pm 0.38$ | $14.9 \pm 4.1$ | $0.033 \pm 0.0018$ | $1.54 \pm 0.36$ | $21.4 \pm 5.4$ | $0.018 \pm 0.001$ | $0.72 \pm 0.1$ | $26.0 \pm 3.7$ | ND | ND | ND |
| Myristyl | $0.022 \pm 0.001$ | $1.74 \pm 0.44$ | $12.6 \pm 3.2$ | $0.030 \pm 0.0016$ | $1.48 \pm 0.35$ | $20.3 \pm 4.9$ | $0.016 \pm 0.001$ | $0.68 \pm 0.07$ | $27.8 \pm 3.76$ | ND | ND | ND |
| Palmytyl | $0.019 \pm 0.0006$ | $1.13 \pm 0.18$ | $16.8 \pm 2.3$ | $0.03 \pm 0.0016$ | $1.13 \pm 0.28$ | $26.5 \pm 6.7$ | $0.015 \pm 0.001$ | $0.56 \pm 0.05$ | $27.2 \pm 3.41$ | ND | ND | ND |

**Figure 7.**
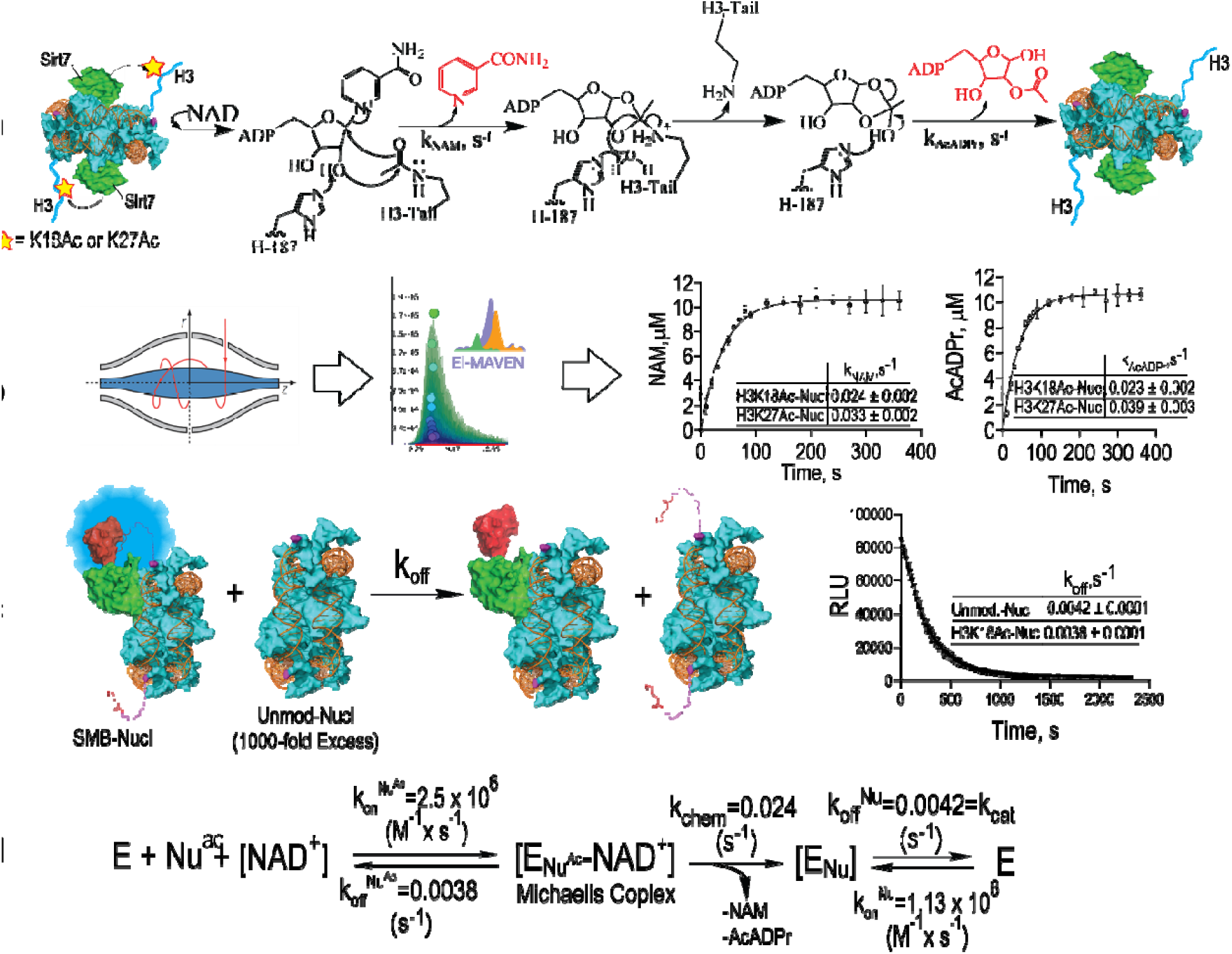
Intramolecular Sirt7 nucleosome deacetylation. Experimental design (a) and LC/MS analysis (b) of the single turnover intramolecular Sirt7 nucleosome deacetylation process. LC/MS-monitored products highlighted in red were quantitated using ElMaven and plotted vs time to determine the corresponding kinetic parameters for H3K18 and H3K27 intramolecular nucleosome deacetylation. c. Experimental design for determination of Sirt7-nucleosome dissociation rate constants (k_off_) using NanoLuc complementation assay. Dissociation rates for H3K18Ac (substrate) and unmodified (product) nucleosomes were determined in the presence of ADPr. d. The enzymatic mechanism of intramolecular Sirt7 nucleosome deacetylation. Individual microscopic rate constants were determined for each step. a).

### Conclusion

In this work we present a comprehensive *in vitro* biochemical analysis of Sirt7 aimed to explain its biological functions. We conducted a detailed systematic investigation of Sirt7 enzymatic activation by DNA, tRNA, and nucleosomes using a broad spectrum of short- and long-chain acylated synthetic peptides, recombinant acetylated nucleosomes, and native chromatin. In the course of this study, we developed several novel biochemical tools and established a strong experimental foundation for prospective sirtuin-specific drug development research. Utilizing the newly developed methodologies, we demonstrated that in contrast to the previous reports^19,22^, tRNA has no significant advantage over DNA as both modulators equally well stimulate Sirt7’s long-chain deacylation activity. More importantly, we identified that Sirt7 preferentially interacts with and is potently activated by nucleosomes towards biologically more abundant acetylated substrates. Sirt7 activation by nucleic acids (DNA and tRNA) is equally effective only towards a narrow range of hydrophobic long-chain acylated substrates while only interaction with nucleosomes leads to potent activation of Sirt7 deacetylation activity consistently with previous biological observations^38^. The direct competition between nucleic acids and nucleosomes further diminishes the biological significance of DNA and tRNA as natural Sirt7 activators in the cell considering the NLS-containing enzyme is localized in the nuclear chromatin-rich environment. For this reason, all prospective studies aimed at developing novel Sirt7-specific inhibitors should be focused on its active physiologically relevant complex with nucleosomes. However, outside of the nuclear envelope, all double-stranded nucleic acids as small as 18-20bp may play an important role as activators of Sirt7 long-chain deacylation activity. Using endogenous chromatin as well as recombinant acetylated substrate nucleosomes, we showed that the site-specifically acetylated nucleosomes act as self-activating substrates for Sirt7. The catalytic efficiency of such an intramolecular deacetylation process is higher compared to intermolecular peptide deacetylation. However, the nucleosome-activated Sirt7 is a highly capable both intermolecular and intramolecular deacetylase and the most specific and catalytically efficient enzyme in the sirtuin family.

## Materials and methods

### Materials

All cloning reagents were obtained from New England Biolab (Gibson assembly Master Mix, Q5-mutagenesis kit) or Thermo Scientific (Fast Digest restriction enzymes). Reagents and building blocks for the solid phase peptide synthesis (SPPS) were from Anaspec, P3Bio, ChemPep, or Aaptec. The Chem Matrix Rink Amide resin was from Sigma. Solvents for SPPS, HPLC, and LC/MS, buffering agents, and additives were from Sigma or Fisher Scientific. Antibiotics, IPTG, DTT, and protease inhibitors (leupeptin, pepstatin, aprotinin, AEBSF) were from Gold Bio. KRX *E*.*coli* cells, NanoBit, and NanoGlow luciferase assay components were from Promega, and L-rhamnose was from Chem-Implex. tRNA from yeast was purchased from Sigma. Corning 384 Well Low Volume Black Round Bottom Polystyrene NBS Microplates were from Sigma. Anti-Histone H3 (acetyl K18) antibody - ChIP Grade (ab1191) was from Abcam, Anti-Histone H3 (acetyl K9) antibody (ab4441) was from Abcam. Secondary antibodies IRDye 680RD anti-mouse and IRDye 800CW anti-rabbit were from Li-Cor.

### Cloning, expression, and purification of Sirt7

The novel Sirt7 *E*.*coli* expression system and purification protocols are described in Supplementary Information (Supplementary figures S1 and S2).

### Electrophoretic mobility shift assays

20 µLs of 400 nM DNA, tRNA, or nucleosomes were mixed in siliconized plastic tubes with equal volume (40 µLs final) of serially diluted Sirt7 in the buffer containing 10 mM Tris, 50 mM NaCl, 0.5 mM TCEP, pH 8.0, yielding the “primary complex mixtures”. The mixtures were incubated for 30 min (RT) with vigorous shaking, and 20 µLs from each tube were mixed with 5x peptide/NAD^+^ solution containing 10 mM Tris, 50 mM NaCl, 0.5 mM TCEP, pH 8.0, 500 µM H3K18Deca, 5 mM NAD^+^. After 30 min incubation (RT) the reactions were quenched with 5 µLs of 6% (v/v) TFA in water and analyzed by HPLC. Reactions containing serially diluted Sirt7 mixed with nucleosome- and nucleic acid-free reaction buffer in the presence of H3K18Deca and NAD^+^ were used as controls to account for non-activated apoSirt7 reactivity. After 30 min incubation (RT) the control reactions were quenched with 5 µLs of 6% (v/v) TFA in water and analyzed by HPLC. The amounts of product released in apoSirt7-catalyzed (control) reactions were substructed from the product released in corresponding reactions catalyzed by Sirt7 in the presence of DNA, tRNA, or nucleosomes. The values of three independent experiments were plotted and presented as bar plots with S.D. (Figure 1).

The remaining 20 µLs of the “primary complex mix” were supplemented with 5 µLs of 50% (w/v) glycerol in 10 mM Tris, 50 mM NaCl, 0.5 mM TCEP, pH 8.0 buffer, and 5 µLs were loaded and resolved on ice using 0.2xTBE 10% acrylamide: bis-acrylamide (59:1) gel for 30 min at 150 volts. A DNA ladder (1 µL of GeneRuler 1 kb or GeneRuler 1 kb Plus; Thermo Fisher) was typically included in the first lane. The gel was stained with SYBR Safe for DNA or SYBR GREEN II (Thermo Fisher) for tRNA and imaged on a Typhoon 9000 imager (GE Healthcare) using Typhoon Control Software (v1.1). SYBR fluorescence was excited at 473 nm and scanned with a 510 nm filter cutoff. For activity assay, the remaining 20 µLs of each mixture

### Fluorescence polarization binding assay

10 µLs of 10 nM FAM-labeled fluorescent double-stranded DNA or LANA-FAM peptide probe were mixed with 10 µLs of serially diluted Sirt7 or nucleosomes in non-binding 384 well low-volume non-binding black plates. Reaction buffer for the al FP experiment contained 10 mM Tris, pH 8.0, 50 mM NaCl, 0.5 mM EDTA. The plates were spun, sealed, and incubated for 5 minutes at 37°C before reading in the BioTek Synergy plate reader using 485/20 nM and 520/10 nM filters for excitation and emission, respectively. Fluorescence polarization values were automatically calculated and the values from at least three independent experiments were plotted with error bars representing S.D. The data were fitted to a specific binding variable Hill slope model embedded in the GraphPad Prism version 8.0.1 package for Windows, GraphPad Software, San Diego, California USA.

### Fluorescence polarization competition assay

Sirt7 (120nM final) and FAM-DNA (10 nM final) were pre-mixed and 10 µLs of the complex were added to 10 µLs of serially diluted (0.2-2000 nM) competitor (tRNA or nucleosomes) in non-binding 384-well low-volume black plates. The plates were spun, sealed, and incubated for 30 minutes at 37°C before reading in the BioTek Synergy plate reader using 485/20 nM and 520/10 nM filters for excitation and emission, respectively. Fluorescence polarization values were automatically calculated, and at least three independent experiments were performed for each condition. The values were plotted and fitted to the variable slope competition model embedded in the GraphPad Prism version 8.0.1 package for Windows, GraphPad Software, San Diego, California USA.

### Sirt7 Michaelis-Menten kinetics with variable NAD^+^ substrate

Serial dilutions of NAD^+^ substrate (10-10000 µM) were prepared in the buffer containing 50 mM Tris, pH 8.0. Concentrated 5x Sirt7 complexes with DNA (2.5 µM Sirt7), tRNA (2.5 µM Sirt7), or nucleosomes (250 nM Sirt7) were prepared as described above. 20 µLs of H3K18Ac- DNB (500 µM) or H3K18Deca-DNB (500 µM) peptide substrates in 50 mM Tris, pH 8.0, 20% DMSO, 1 mM DTT were mixed with 20 µLs of serially diluted NAD^+^. The substrate mixture was pre-incubated at 37°C for 10 minutes, and the reaction was initiated with 10 µLs of each Sirt7 complex. Reactions were incubated for 30-60 minutes, quenched with 10 µLs of 6% TFA, and analyzed by HPLC as previously described. The data was plotted and fitted to the Michaelis Menten equation using GraphPad Prism.

### Sirt7 Michaelis-Menten kinetics with a variable peptide substrate

The preparation of synthetic peptides is described in the supplementary information. Serial dilutions of synthetic acylated substrate peptides were prepared in the buffer containing 50 mM Tris, pH 8.0, 20% DMSO. For nucleic acid activation, 2 µLs of 1 mM DNA or tRNA were added to 1 mL of 1 µM Sirt7 solution in 50 mM Tris, pH 8.0. The resulting complex solution was supplemented with 2 µM nicotinamidase and equilibrated at 37°C with vigorous 1200 rpm shaking for 15 minutes. For nucleosome activation assay 200 nM Sirt7 solution in 50 mM Tris, pH 8.0, was mixed with concentrated stock nucleosomes (10-25 µM, 400 nM final). The resulting complex was adjusted to 100 nM Sirt7 with 50 mM Tris, pH 8.0 buffer, supplemented with 2 µM nicotinamidase, and equilibrated at 37°C with vigorous 1200 rpm shaking for 15 minutes. Thus, all complexes were prepared with a two-fold stoichiometric excess of nucleic acid or nucleosomes over Sirt7. 20 µLs of each complex were mixed with 20 µLs of the serially diluted peptide at 37°C, and the reaction was initiated by the addition of 10 µLs of 10 mM freshly prepared buffered NAD^+^ co-substrate in 50 mM Tris, pH 8.0. Reactions were incubated for 5-60 minutes at 37°C and quenched with 10 µLs of 6% TFA. Quenched reactions were analyzed by HPLC equipped with a C18 Kinetex column (100 Å, 100 × 4.6 mm, 2.6 μm, Phenomenex). Elution of substrate and the deacetylated product was monitored at 214 nm, 275 nm, or 400 nm. For tryptophan-labeled peptides (Figure 2) at least a 20-30 minute linear acetonitrile gradient at 1.6 mL min^−1^ was necessary with continuous monitoring at 214 or 280 nm. For DNB-labeled peptides reactions were monitored at 400nm and the length of the acetonitrile gradient was reduced to 6 min at 1.6 mL min^−1^. The deacetylated and acetylated peptide peaks were integrated, and the ratio was multiplied by the total substrate concentration to calculate the product concentration. Three independent experiments were conducted for each run and the data was plotted and fitted to the Michaelis Menten equation using GraphPad Prism.

### Sirt7 kinetic titration assay

Serial dilution of DNA (1-1000 µM), tRNA(1-1000 µM) or nucleosomes (0.1-100 nM) were prepared in 50 mM Tris, pH 8.0. 20 µLs of the master mix containing 20 nM Sirt7, 50 µM H3K18Deca peptide, and 5 mM NAD^+^ in 50 mM Tris, pH 8.0 buffer were added to 20 µLs of serially diluted modulators (nucleic acids or nucleosomes) in siliconized PCR strip tubes. Reactions were incubated at 37°C for 5-8 hours in the BioRad thermal cycler set to infinite isothermal incubation with 50°C heated lid to prevent evaporation. Reactions were quenched with 10 µLs of 6% TFA and analyzed by HPLC as previously described. The values were fitted to a specific binding variable Hill slope model embedded in the GraphPad Prism version 8.0.1 package for Windows, GraphPad Software, San Diego, California USA.

### Singe turnover nucleosome deacetylation assay

The preparation of recombinant acetylated nucleosomes is described in the supplementary information. 200 µLs of H3K18Ac or H3K27Ac nucleosomes (6 µM) in 50 mM Tris, pH 8.0, 1 mM DTT were mixed with 4 µLs of 512 µM stock Sirt7 (10 µM final), and the complex was equilibrated with vigorous shaking (1200 rpm) for 30 minutes at 37°C. To 20 µLs of buffered freshly prepared 10 mM NAD^+^ (10x) solution pre-incubated at 37°C, 180 µLs of the Sirt7- nucleosome complex were added with vigorously mixing. Every 20 seconds 10 µLs of mixed reaction were quenched by 10 µL of 1% (v/v) mass-spectrometry grade formic acid in water. For the control experiment, 20 µLs of the same 10 mM NAD^+^ (10x) solution at 37°C were mixed with 180 µLs of reaction buffer without Sirt7. Reactions were analyzed by Thermo Scientific Q Exactive Orbitrap mass spectrometer coupled to a Vanquish UHPLC and equipped with a Vydac 201SP104 C18 column using 3-95% 20 min water-acetonitrile (0.1% formic acid) gradient at 0.5 ml/min. Nicotinamide and acetyl-ADP-ribose product peaks were integrated and quantified using standard calibration curves. The first-order rate constants (k) of intramolecular nucleosome deacetylation were determined by fitting quantified product to a single-exponential equation, P = [S]_0_ (1 − e^−kt^), where P is the concentration of product formed, [S]_0_ is the initial concentration of total nucleosome acetylation sites.

### NanoBit nucleosome equilibrium binding assay

The details of cloning, expression, and purification of NanoBit assay components are described in the supplementary information. Serial dilutions of small bit nucleosomes (SMB-Nucs, 0.5-50 nM) were prepared in the NanoLuc assay buffer containing 50 mM Tris, pH 8.0, 250 µg/ml bovine serum albumin (BSA). In 384 well low-volume non-binding black plates 9 µLs of Sirt7- CLB (250 pM) or Sirt7-CLB+2mM ADPr in the same buffer were mixed with 9 µLs of serially diluted SMB-Nucs or SMB-AcNucs. The plates were spun (500 x g) for 30 seconds, sealed, and incubated for 20 minutes at room temperature. Following this incubation, 2 µL of 20% (v/v) NanoGlow luciferase substrate solution in the NanoLuc assay buffer (without BSA) were (final dilution factor was 50x from the supplied Promega stock solution cat#N1110). The plate was sealed and inserted into the BioTek Synergy plate reader. Reactions were equilibrated at 37°C for 5 minutes and read continuously for 15 minutes (1 mm probe offset, sensitivity 100-120, 0.5 second integration) every 30 seconds to control for steady-state signal stability. At least three independent experiments were conducted, and the averaged relative luciferase unit (RLU) values were plotted vs nucleosome concentration. Dissociation constants (K_d_) were determined by fitting data to a one site-specific binding model using GraphPad Prism.

### NanoBit Sirt7-nucleosome complex dissociation assay

10 µLs of Sirt7-CLB (400 pM) in the NanoLuc assay buffer (50 mM Tris, pH 8.0, 250 µg/ml BSA) were mixed with 10 µLs SMB-Nucs (6 nM) or SMB-AcNucs in the same buffer with additional 3 mM ADPr (1.5 mM in the 20 µL final volume). The complex was equilibrated at 37° for 20 minutes and mixed with 1 µL of NanoGlow luciferase substrate (Promega cat#N1110). The mixture was pre-incubated at 37°C for 5 min with vigorous shaking (1200 rpm), and 19 µLs of the complex were added to 1 µL of untagged 10x stock competitor nucleosomes (25 µM stock, 2.5 µM final) in the 384 black low volume non-binding plate. The plate was immediately sealed and inserted into the BioTek Synergy plate reader pre-equilibrated at 37°C. RLU values were measured (1 mm probe offset, sensitivity 100- 120, 0.5 second integration) with 5 s time intervals over 1 hour. The control experiment was carried out in the absence of competing nucleosomes to ensure that complemented luciferase signal remains stable during the experiment. The delay period from the addition of competitor nucleosome to the first BioTek Synergy read (15-20 s) was accurately recorded and accounted for in the final plotting. The RLU values from three independent experiments were plotted and fitted to one phase dissociation exponential decay model using GraphPad Prism to determine dissociation k_off_ rates for acetylated and unmodified nucleosomes.

### NanoBit equilibrium competition assay

16 µLs of pre-equilibrated Sirt7-CLB (3 nM) and SMB-Nucs (300 pM) were mixed with 3 µLs of serially diluted (0.001-70 µM) competitor DNA or tRNA in the NanoLuc assay buffer in the 384 black low volume non-binding plate. The plate was sealed and equilibrated for 20 minutes at 37°C. The equilibrated reactions were spun (500xg) and 1 µL of freshly prepared 50%(v/v) NanoGlow luciferase substrate (Promega cat#N1110) was added to each well. The plate was immediately sealed and inserted into the BioTek Synergy plate reader. The luminescence was continuously monitored (1 mm probe offset, sensitivity 100-120, 0.5 second integration) with 10 seconds time intervals over 20 minutes at 37°C. A stable signal was achieved after 5-10 minutes and the averaged RLU values from three independent experiments were plotted and fitted to the variable slope competition model using GraphPad Prism.

### Chromatin purification and western dot-blot analysis

PC3 cancer cell lines grown to 60% confluency in high glucose DMEM media were treated with HDAC inhibitor mix (10 μM SAHA, 10 mM sodium butyrate, and 10 mM nicotinamide) in the same media for 24 hours before harvesting. Chromatin was isolated, purified, and digested to native oligonucleosomes as previously described.^35^ Deacetylation reaction mixture (200µLs) containing 10 µM native oligonucleosomes, 1 mM NAD^+^, 1 µM nicotinamidase, and 1 µM of Sirt2, Sirt6, or Sirt7, was incubated for 2 hours at 37°C with vigorous (1000 rpm) shaking. Reactions were quenched and analyzed by Thermo Q Exactive Orbitrap mass spectrometer coupled to a Dionex Ultimate 3000 RSLC nano UPLC with a Waters Atlantis reverse phase column (100 μm × 150 mm) as previously described.^35^During the incubation period small 4 µLs aliquots were quenched by mixing with 400 µLs of 1% TFA at 0, 5, 10, 15, 30, 60, and 120 minutes. 100 µLs from each reaction were adjusted to 1 ml with DI water and loaded on nitrocellulose membrane using Bio-Rad Bio-Dot 96 well microfiltration blotting device per manufacturer instructions. The membrane was blocked with 5% BSA in TBST and blotted overnight at 4°C with a mixture of anti-H3(mouse) total and H3K9Ac or H3K27Ac (rabbit) antibodies in the same buffer. After extensive washing, a mixture of secondary antibodies (IRDye 680RDandi-mouse and IRDye 800CW anti-rabbit, Li-Cor, 5000x dilution in the same buffer) was added and incubated for 1 hour at room temperature. After 3 final washes with TBS to remove Tween-20 detergent the membranes were imaged using LI-COR Odyssey imager. Individual spots were quantitated using Li-Cor ImageStudio software.

## Acknowledgments

We want to thank Prof. Peter Lewis for donating bacterial plasmids for the expression and purification of human histones. This work was supported by NIH grant R01 GM65386.

## Author contributions

V.I.K. designed the study and performed cloning, protein expression, purification, kinetic /binding assays, and data analysis. V.I.K. and M.A.K. synthesized peptides, designed, optimized, and validated NanoLuc binding and dissociation experiments; V.I.K. and W.H.L. designed and conducted EMSA; V.I.K. performed mammalian tissue culture experiments, chromatin isolation, and LC/MS analysis. V.I.K. and J.M.D. wrote the original manuscript draft; V.I.K., W.H.L., M. A. K., and J.M.D. proofread and edited the manuscript.

