## Supporting Information for "Potent activation of NAD^+^-dependent deacetylase Sirt7 by nucleosome binding"

**MATERIALS AND METHODS**

**Cloning, expression, and purification of Sirt7.** The primary protein sequence of Sirt7 (Uniprot Q9NRC8) was reverse translated using OPTIMIZER,^1^ and the expression cassette was synthesized by IDT. The codon-optimized G-block expression cassette containing rhamnose promoter upstream of Sirt7-encoding sequence was cloned into the backbone of the pet3d (Novagen) vector pre-digested with Bgl-II and BamHI restriction enzymes yielding pet3d-Sirt7 construct (Supplementary Information). The protein was expressed in KRX *E.coli* cells (Promega) that were grown at 37^o^C in 2xYT media until the optical density (OD_600_) reached 1.2-1.5. To induce Sir7 expression the cultures were supplemented with 10 milliliters of 10% (w/v) rhamnose solution (final rhamnose concertation 0.1% (w/v)). The protein expression continued at room temperature for 16-18 hours, and cells were harvested by centrifugation. *E.coli* pellet was re-suspended (30% (w/v)) in the Ni binding buffer containing 50 mM Tris, pH 8.0. 25mM Imidazole, 1M NaCl, 1 mM β-mercaptoethanol (βME), 1 M NaCl, supplemented with protease inhibitors (leupeptin, pepstatin, AEBSF) and lysozyme. The cells were sonicated on ice in a beaker equipped with a magnetic stirring bar (40% amplitude, 10 second pulse ON/59 seconds OFF) for 30 minutes, and 10%(w/v) polyethyleneimine (M.w. 50000) solution (pH 8)^2^ was directly added to the lysed cells with stirring to the final concentration of 0.5% (w/v). The viscous solution containing lysed cell debris and precipitated DNA was centrifuged, and solid ammonium sulfate was added to the clear supernatant to 40-45% saturation (calculated at 4^o^C). After 10 minutes of vigorous stirring precipitated Sirt7 was centrifuged and re-suspended in 20 milliliters of the Ni binding buffer, and loaded at 2ml/min flow rate on 5 ml Ni-chelating column using AKTA FPLC system (GE). The resin was extensively washed with the Ni loading buffer (without protease inhibitors) followed by 100 milliliters of Wash Buffer#1 containing 10mM Tris, 1mM βME, and finally with 200 milliliters of Wash Buffer#2 containing 10 mM Tris-Base, 150 mM imidazole, 80 mM NaCl, pH 8.0. All washing steps with Ni loading buffer, wash buffer #1 and wash buffer #2 were conducted using the same pump A of the AKTA FPLC. After the last wash#2 the protein was eluted with a 120 milliliters linear gradient using Ni elution buffer containing 50 mM Tris, 300 mM imidazole, 1M NaCl, 1 mM βME, pH 8.0. Purified Sirt7 was analyzed by SDS-PAGE (Figure S1, b), and all eluted protein was dialyzed in TEV cleavage buffer containing 50 mM HEPES, pH 7.5, 500 mM NaCl, 1 mM DTT, and recombinant TEV-protease was added to cleave the N-terminal hexa-histidine tag overnight at 4^o^C. Cleaved Sirt7 was diluted two-fold to adjust NaCl concertation down to 250 mM with SP-loading buffer containing 50 mM HEPES, pH 7.5, 1 mM DTT, and loaded on SP-HP column using AKTA FPLC. After extensive wash with SP-loading buffer the protein was eluted with a 150 milliliter linear gradient against 50 mM HEPES, pH 7.5, 1M NaCl, 1 mM DTT. Purified Sirt7 was analyzed by SDS-PAGE (Figure S1, b), fractions containing Sirt7 were concentrated to 7-10 mg/ml. Small 10 µL aliquots were flash-frozen in the liquid nitrogen using 10% (v/v) glycerol as a cryoprotector.

>Optimized pet3d(B-B)-Sirt7WT (this work) **CAI Value: 0.70**

agatctCACCACAATTCAGCAAATTGTGAACATCATCACGTTCATCTTTCCCTGGTTGCCAATGGCCCATTTTCCTGTCAGTAACGAGAAGGTCGCGAATTCAGGCGCTTTTTAGACTGGTCGTAGGGAGACCACAACGGTTTCCCTCTAGAAATAATTTTGTTTAACTATAAGA**AGGAGA**TATACAT**ATGCATCATCATCATCATCACAGCGGTGGGGAAGATTTGTATTTTCAG**GCAGCGGGGGGTTTAAGCCGTTCGGAGCGCAAAGCTGCAGAACGTGTGCGCCGTTTACGTGAGGAGCAGCAGCGTGAACGTTTACGCCAGGTATCGCGTATTTTGCGCAAAGCAGCTGCAGAACGTAGCGCAGAAGAGGGGCGTTTGCTGGCTGAAAGCGCGGATCTGGTTACAGAGCTGCAGGGGCGTAGTCGCCGCCGCGAAGGGCTGAAACGTCGCCAGGAAGAGGTGGTAGACGATCCGGAAGAATTGCGTGGGAAAGTCCGCGAGTTGGCGTCGGCTGTTCGTAACGCGAAATATTTGGTGGTATACACAGGGGCAGGTATCTCGACGGCAGCGAGCATCCCTGACTATCGTGGGCCTAATGGGGTGTGGACGTTACTGCAGAAAGGCCGTAGTGTGTCGGCGGCTGACTTGAGCGAAGCGGAACCTACTCTGACGCATATGAGTATCACGCGTCTGCACGAGCAGAAGCTGGTTCAGCACGTGGTAAGCCAGAATGTGGATGGGCTGCACTTACGTAGCGGGTTGCCTCGCACAGCTATCTCGGAGTTGCATGGGAACATGTATATCGAAGTCTGCACTAGTTGTGTCCCTAATCGTGAGTATGTACGTGTTTTTGATGTCACTGAGCGCACGGCATTGCATCGTCACCAGACGGGTCGCACGTGCCATAAATGCGGGACGCAGTTACGCGACACTATTGTGCATTTCGGCGAGCGTGGTACTTTAGGGCAGCCTTTAAATTGGGAAGCAGCTACGGAAGCGGCGTCGCGTGCGGACACTATCCTGGTCCTGGGGAGTAGCCTGAAAGTGCTGAAAAAGTATCCTCGTTTGTGGTGCATGACGAAACCGCCTTCGCGTCGCCCTAAACTGTATATTGTGAACTTGCAGTGGACGCCTAAGGACGACTGGGCAGCGTTAAAACTGCACGGCAAGGTGGACGATGTGATGCGCTTGCTGATGGCGGAATTAGGGTTGGAAATTCCGGCATATAGCCGTTGGCAGGACCCGATCTTTAGTTTGGCGACACCTCTGCGTGCTGGTGAAGAAGGTAGTCATTCGCGTAAGAGCCTGGTGCGCAGCCGTGAAGAAGCGCCACCTGGGGATCGTGGTGCGCCTCTGAGCAGTGCGCCGATCTTGGGTGGGTGGTTCGGTCGTGGGGTGACGAAACGCACTAAACGTAAAAAAGTTACG**TAA**ggatccGGCTGCTAACAAAGCCCGAAAGGAAGCTGAGTTGGCTGCTGCCACCGCTGAGCAATAA

Cyan – rhamnose promoter, Blue - N-terminal 6his tag, Red- TEV protease cleavage site, Green – codon-optimized human Sirt7, represent N-and C-terminal deletions that were prepared in addition to the full length (FL) Sirt7.

>Sirt7ΔCys. Translated sequence.

MHHHHHHSGGEDLYFQAAGGLSRSERKAAERVRRLREEQQRERLRQVSRILRKAAAERSAEEGRLLAESADLVTELQGRSRRREGLKRRQEEVVDDPEELRGKVRELASAVRNAKYLVVYTGAGISTAASIPDYRGPNGVWTLLQKGRSVSAADLSEAEPTLTHMSITRLHEQKLVQHVVSQNVDGLHLRSGLPRTAISELHGNMYIEVCTSCVPNREYVRVFDVTERTALHRHQTGRTCHKCGTQLRDTIVHFGERGTLGQPLNWEAATEAASRADTILVLGSSLKVLKKYPRLWCMTKPPSRRPKLYIVNLQWTPKDDWAALKLHGKVDDVMRLLMAELGLEIPAYSRWQDPIFSLATPLRAGEEGSHSRKSLVRSREEAPPGDRGAPLSSAPILGGWFGRGVTKRTKRKKVT

Purple: cysteines substituted with V to improve oxidative stability of the enzyme.

**Cloning, expression, and purification of Sirt7-CLB for NanoLuc assay.** The DNA fragment of the C-terminal Large Bit (CLB) component was amplified from pBiT1.1-C vector (Promega) while the codon-optimized Sirt7 sequence was amplified from the previously described pet3d-Sirt7 plasmid. The fragments were ligated with pet14b vector pre-digested with NdeI-XhoI restriction endonucleases using HiFi Gibson assembly Master Mix (NEB). The protein was purified as previously described for pet3d-Sirt7 with an additional Superdex 16/60 size-exclusion purification step. After SP-HP ion exchange chromatography the concentrated protein was loaded on Superdex 200 (16/600) pre-equilibrated with 50 mM Tris, pH 8.0, 2M NaCl, 1 mM DTT (Figure S1, e, f). Homogeneous fractions were pulled together, concentrated to 1 ml, and flash-frozen in liquid nitrogen using 10% glycerol as cryoprotecting agent.

>pet14b-St7-CLB Green – Sirt7WT sequence; Purple – Large Bit (CLB) fragment of the NanoLuc split luciferase complementation system; Red – flexible linker.

CCATGGGCAGCAGCCATCATCATCATCATCACAGCAGCGGCCTGGTGCCGCGCGGCAGCCAAGGTTTAAGCCGTTCGGAGCGCAAAGCTGCAGAACGTGTGCGCCGTTTACGTGAGGAGCAGCAGCGTGAACGTTTACGCCAGGTATCGCGTATTTTGCGCAAAGCAGCTGCAGAACGTAGCGCAGAAGAGGGGCGTTTGCTGGCTGAAAGCGCGGATCTGGTTACAGAGCTGCAGGGGCGTAGTCGCCGCCGCGAAGGGCTGAAACGTCGCCAGGAAGAGGTGGTAGACGATCCGGAAGAATTGCGTGGGAAAGTCCGCGAGTTGGCGTCGGCTGTTCGTAACGCGAAATATTTGGTGGTATACACAGGGGCAGGTATCTCGACGGCAGCGAGCATCCCTGACTATCGTGGGCCTAATGGGGTGTGGACGTTACTGCAGAAAGGCCGTAGTGTGTCGGCGGCTGACTTGAGCGAAGCGGAACCTACTCTGACGCATATGAGTATCACGCGTCTGCACGAGCAGAAGCTGGTTCAGCACGTGGTAAGCCAGAATGTGGATGGGCTGCACTTACGTAGCGGGTTGCCTCGCACAGCTATCTCGGAGTTGCATGGGAACATGTATATCGAAGTCTGCACTAGTTGTGTCCCTAATCGTGAGTATGTACGTGTTTTTGATGTCACTGAGCGCACGGCATTGCATCGTCACCAGACGGGTCGCACGTGCCATAAATGCGGGACGCAGTTACGCGACACTATTGTGCATTTCGGCGAGCGTGGTACTTTAGGGCAGCCTTTAAATTGGGAAGCAGCTACGGAAGCGGCGTCGCGTGCGGACACTATCCTGGTCCTGGGGAGTAGCCTGAAAGTGCTGAAAAAGTATCCTCGTTTGTGGTGCATGACGAAACCGCCTTCGCGTCGCCCTAAACTGTATATTGTGAACTTGCAGTGGACGCCTAAGGACGACTGGGCAGCGTTAAAACTGCACGGCAAGGTGGACGATGTGATGCGCTTGCTGATGGCGGAATTAGGGTTGGAAATTCCGGCATATAGCCGTTGGCAGGACCCGATCTTTAGTTTGGCGACACCTCTGCGTGCTGGTGAAGAAGGTAGTCATTCGCGTAAGAGCCTGGTGCGCAGCCGTGAAGAAGCGCCACCTGGGGATCGTGGTGCGCCTCTGAGCAGTGCGCCGATCTTGGGTGGGTGGTTCGGTCGTGGGGTGACGAAACGCACTAAACGTAAAAAAGTTACGCTGCTCGAGGGTGGTGGCGGGAGCGGAGGTGGAGGGTCGTCAGGTGTCTTCACACTCGAAGATTTCGTTGGGGACTGGGAACAGACAGCCGCCTACAACCTGGACCAAGTCCTTGAACAGGGAGGTGTGTCCAGTTTGCTGCAGAATCTCGCCGTGTCCGTAACTCCGATCCAAAGGATTGTCCGGAGCGGTGAAAATGCCCTGAAGATCGACATCCATGTCATCATCCCGTATGAAGGTCTGAGCGCCGACCAAATGGCCCAGATCGAAGAGGTGTTTAAGGTGGTGTACCCTGTGGATGATCATCACTTTAAGGTGATCCTGCCCTATGGCACACTGGTAATCGACGGGGTTACGCCGAACATGCTGAACTATTTCGGACGGCCGTATGAAGGCATCGCCGTGTTCGACGGCAAAAAGATCACTGTAACAGGGACCCTGTGGAACGGCAACAAAATTATCGACGAGCGCCTGATCACCCCCGACGGCTCCATGCTGTTCCGAGTAACCATCAACAGCTAAGGATCCGGCTGCTAACAAAGCCCGAAAGGAAGCTGAGTT

>Sirt7-CLB. Translated Sequence

MGSSHHHHHHSSGLVPRGSQGLSRSERKAAERVRRLREEQQRERLRQVSRILRKAAAERSAEEGRLLAESADLVTELQGRSRRREGLKRRQEEVVDDPEELRGKVRELASAVRNAKYLVVYTGAGISTAASIPDYRGPNGVWTLLQKGRSVSAADLSEAEPTLTHMSITRLHEQKLVQHVVSQNVDGLHLRSGLPRTAISELHGNMYIEVCTSCVPNREYVRVFDVTERTALHRHQTGRTCHKCGTQLRDTIVHFGERGTLGQPLNWEAATEAASRADTILVLGSSLKVLKKYPRLWCMTKPPSRRPKLYIVNLQWTPKDDWAALKLHGKVDDVMRLLMAELGLEIPAYSRWQDPIFSLATPLRAGEEGSHSRKSLVRSREEAPPGDRGAPLSSAPILGGWFGRGVTKRTKRKKVT**LLEGGGGSGGGGSSG**VFTLEDFVGDWEQTAAYNLDQVLEQGGVSSLLQNLAVSVTPIQRIVRSGENALKIDIHVIIPYEGLSADQMAQIEEVFKVVYPVDDHHFKVILPYGTLVIDGVTPNMLNYFGRPYEGIAVFDGKKITVTGTLWNGNKIIDERLITPDGSMLFRVTINS

**Cloning, expression, and purification of SmBit-H2B.** Human H2B sequence (Q5QNW6) was PCR-amplified from pRUTH-H2B.bk vector and cloned into pet3d plasmid containing N-terminal small-bit linker tag and N-terminal TEV-cleavable hexa-histidine tag. The plasmid was transformed into KRX cells (Promega). The cultures were grown in 2xYT and protein expression was induced at OD^600nm^=0.8-1.0 with 0.1% (w/v) rhamnose solution as previously described for pet3d-Sirt7. Expression continued at 37^o^C for 4 hours, and harvested cells were lysed by sonication in the denaturing Ni loading buffer containing 50 mM Tris, pH 8.0. 6 M Guanidinium chloride (GuCl), 1mM βME, 1mM AEBSF. The supernatant was loaded on Ni-chelating column (GE) pre-equilibrated with the same Ni loading buffer. After extensive wash the histones were eluted using 120 milliliter linear gradient against buffer B containing 300 mM Imidazole. Collected fractions were dialyzed twice in 2 liters of SP-loading buffer containing 10 mM Ammonium Acetate, pH 6, and 1mM βME. Dialyzed SMB-H2B was loaded on a joint Q-SP column that was extensively washed prior to elution. Q-FF column was disconnected, and SP-HP column was eluted using 120 milliliter linear gradient against buffer B containing 10 mM ammonium acetate, pH 6.0 and 600 mM NaCl. Eluted histone was dialyzed in 5 liters of 10 mM ammonium acetate, 1mM βME, flash-frozen and lyophilized.

>pet3d-SMB-H2B histone. Yellow-his tag, Green-TEV cleavage site, Cyan-small bit tag, Grey – linker, Purple – human H2B (Q5QNW6).

ATGGGACATCACCACCACCACCATGGAGAGAACTTGTACTTCCAGGGTGTGACTGGCTACCGCTTATTTGAGGAGATCTTAGGGTCTTCTGGTGGAGGAGGGAGCGGTGGGGGAGGCTCTTCTGGAGGAGCTCCTGACCCCGCCAAATCTGCACCGGCTCCAAAAAAGGGGTCGAAGAAAGCAGTGACGAAGGTTCAAAAAAAGGACGGGAAAAAACGTAAACGCTCCCGTAAAGAAAGCTACAGCGTATATGTGTATAAGGTGTTGAAACAAGTCCATCCAGATACTGGTATCTCATCTAAGGCGATGGGCATTATGAACAGTTTTGTTAACGACATTTTTGAGCGTATTGCGGGGGAAGCTAGCCGCCTGGCGCACTATAATAAACGTAGTACAATCACTTCACGTGAAATCCAAACTGCTGTTCGTTTATTACTGCCAGGAGAGTTAGCCAAACACGCCGTAAGTGAAGGAACCAAGGCGGTTACCAAGTACACCTCTGCAAAGTAA

>Translated. SMB-H2B

MGHHHHHHGENLYFQGVTGYRLFEEILGSSGGGGSGGGGSSGGAPDPAKSAPAPKKGSKKAVTKVQKKDGKKRKRSRKESYSVYVYKVLKQVHPDTGISSKAMGIMNSFVNDIFERIAGEASRLAHYNKRSTITSREIQTAVRLLLPGELAKHAVSEGTKAVTKYTSAK-

**Preparation of nucleosome core particles.** 6his-Tev-H2A, 6his-Tev -H2B, 6his-Tev -H3, and 6his-Tev-H4 histones were re-suspended in the buffer containing 20 mM HEPES pH 7.5, 6M GuHCl, 1 mM EDTA, 1 mM DTT to the final total concentration of 1 mg/ml (0.25 mg/ml for each individual histone). Solution was dialyzed overnight against 5L of refolding buffer containing 20 mM HEPES pH 7.5, 1 mM EDTA, 2M NaCl, 1 mM βME. Precipitated aggregates were separated by centrifugation, and the octamers were concentrated to 1 mL and purified over Superdex200 (16/600) size-exclusion column pre-equilibrated with fresh refolding buffer. Octamer containing fractions were pulled together and 6his-TEV protease was added to cleave hexa-histidine tags from the constituent histones overnight at 4^o^C. Cleaved octamers were concentrated to 1 ml and purified over Superdex200 (16/600) size-exclusion column. The final concentration of cleaved human octamers was determined using combined extinction coefficient ɛ_280_ = 44700 M^-1^cm^-1^. SmBt-H2B and acK-containing histones were used instead of corresponding canonical histones to produce octamers for NanoLuc and Sirt7 nucleosome deacetylation assays, respectively. Refolded octamers were adjusted to 2 µM using 20 mM HEPES pH 7.5, 1 mM EDTA, 2 M NaCl, 1 mM βME, mixed with equal volume of 2 µM Widom 601 DNA prepared as previously descried^3^ in the same buffer, and dialyzed overnight. To assemble nucleosome core particles low ionic strength buffer containing 10 mM TRIS pH 8.0, 1 mM EDTA, 1 mM βME was titrated to dialyzing nucleosomes overnight to reach the final NaCl concentration of 50 mM. The quality of the assembled nucleosome core particles was evaluated using native electrophoretic mobility shift assays.

**Expression and purification of canonical histones:** pRUTH-H2A.a4 (kanR), pRUTH-H2B.bk (kanR), pRUTH-H3.2 C110A (kanR), pYokoTEVH4 (AmpR) vectors were obtained from Dr. Peter Lewis, PhD. All plasmids were transformed into KRX *E.coli* (Promega). The cells were cultured in 2xYT media until OD^600nm^=0.8-1.0 and protein expression was induced with 1 mM IPTG, 0.1% (w/v) rhamnose at 37^o^C for 5 hours (2 hours for H4). All histones were purified using protocol described for SmBt-H2B. Purified histones were dialyzed in 10 mM ammonium acetate, 1 mM βME, and concentration was determined using calculated extinction coefficients ɛ_280_=5960 M^-1^cm^-1^(6hisH2A), ɛ_280_= 7450 M^-1^cm^-1^ (6hisH2B), ɛ_280_= 5960 M^-1^cm^-1^ (6hisH3), and ɛ_280_=7450 M^-1^cm^-1^ (6hisH4).

**Expression and purification of acetylated histones:** Amber codon was site-specifically introduced into pRUTH-H3.2-C110A (kanR) vector yielding pRUTH-H3.2 C110A, K18acK and pRUTH-H3.2-C110A, K27acK plasmids. Acetylated histones were produced in cobB deficient strain of *E.coli* in 2xYT media supplemented with 10 mM acetylated lysine amino acid, as previously described.^4^

**Activating dsDNA:** The self-complementary DNA oligomers (Table S1) were synthesized by IDT and re-suspended in the alignment buffer containing 10 mM Tris, 150 mM NaCl, 1 mM EDTA. Solution was heated up to 98^o^ C and gradually cooled down to 23^o^C (room temperature) over the period of 2 hours. Re-aligned double-stranded DNAs were additionally purified using preparative HPLC,^5^ lyophilized, and re-suspended in 10 mM Tris, 150 mM NaCl, 1 mM EDTA. Concertation was determined using NanoDrop and IDT-provided extinction coefficients.

**Solid phase peptide synthesis.** All peptides were manually synthesized on Chem-Matrix Rink-Amide resin (Sigma) using DIC/OXYMA^6^ mediated coupling of 4x molar excess of Fmoc-protected amino acids in N-methyl pyrrolidone, followed by chloranil test, acetic anhydride capping and 20% piperidine in DMF deprotection of the N-terminal Fmoc. The N-terminus of tryptophan containing peptides W-GKAQTARK*STGGK was not capped to improve peptide solubility. For peptides synthesized without N-terminal tryptophan a 10 fold excess of 1,4-dinitrobenzene fluoride/DIEA (Sanger reagent) in DMF was used to make DNB-GKAQTARK*STGGK peptides. For FAM labeling of SPPS assembled LANA probe FAM-GMRLRSGRSTGAPLTRGS the final capping step was substitutes with carboxy-fluorescein coupling using HATU as a coupling agent**.** For the acylated substrate peptides the Fmoc-Lys(ivDDE)-OH was incorporated into the sequence GKAQTAR**K***STGGK to allow post-synthetic global acylation with acid anhydrides. The ivDDE-protecting group was removed using 3x 2% (v/v) hydrazine/DMF treatment, the resin was extensively washed with dichloromethane and dried under vacuum to increase the efficiency of the following global modification.^7.^ Selectively deprotected lysine peptide was treated with 0.1-0.5M symmetrical acid anhydrides (acetic, butyric, hexanoic, octanoic, decanoic) in DMF or hydrophobic acid chlorides (dodecanoic, myristic, palmitic, stearic) in dichloromethane using DIEA as a base. The process was monitored using chloranil test and repeated if necessary. Peptides were cleaved with TFA (Reagent K), precipitated with diethyl ether, and C18-HPLC-purified to achieve 98%+ purity.

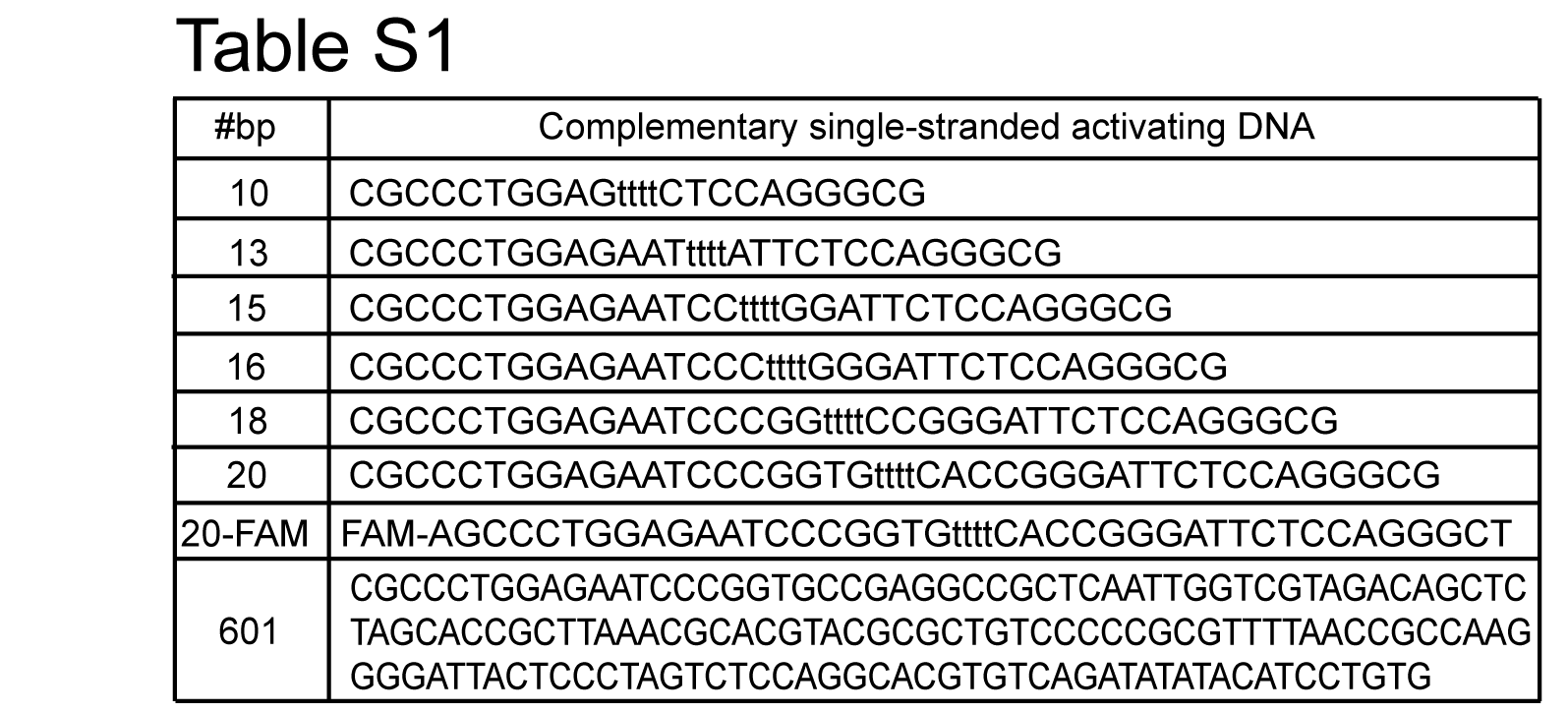
Table S1. Activating DNA sequences used for Sirt7 activation.

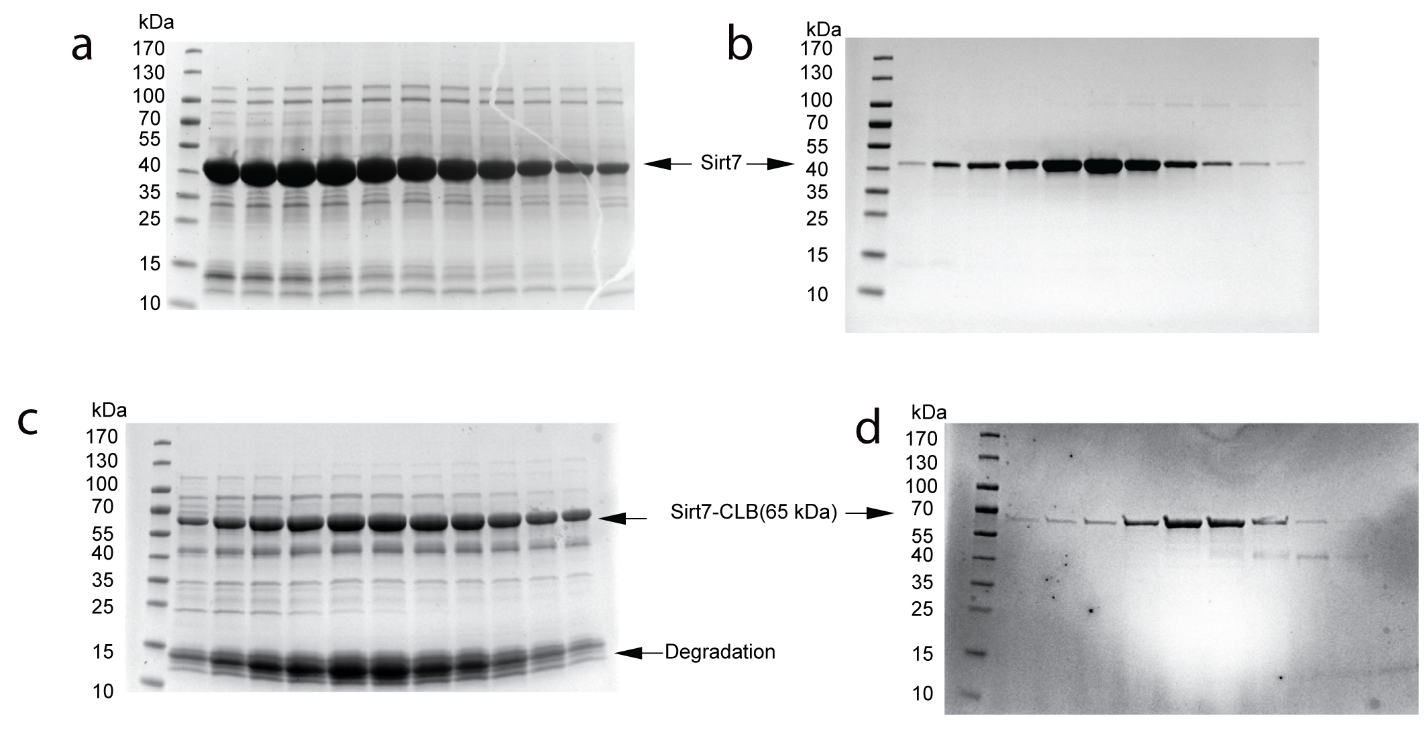

**Supporting figure S1.** SDS-PAGE analysis of Sirt7 purification. Sirt7WT fractions eluted from Ni-chelating (a) and SP-HP (b) cation exchange column. c. Sirt7-CLB fractions eluted from Ni-chelating and d Superdex 200 (d) columns.

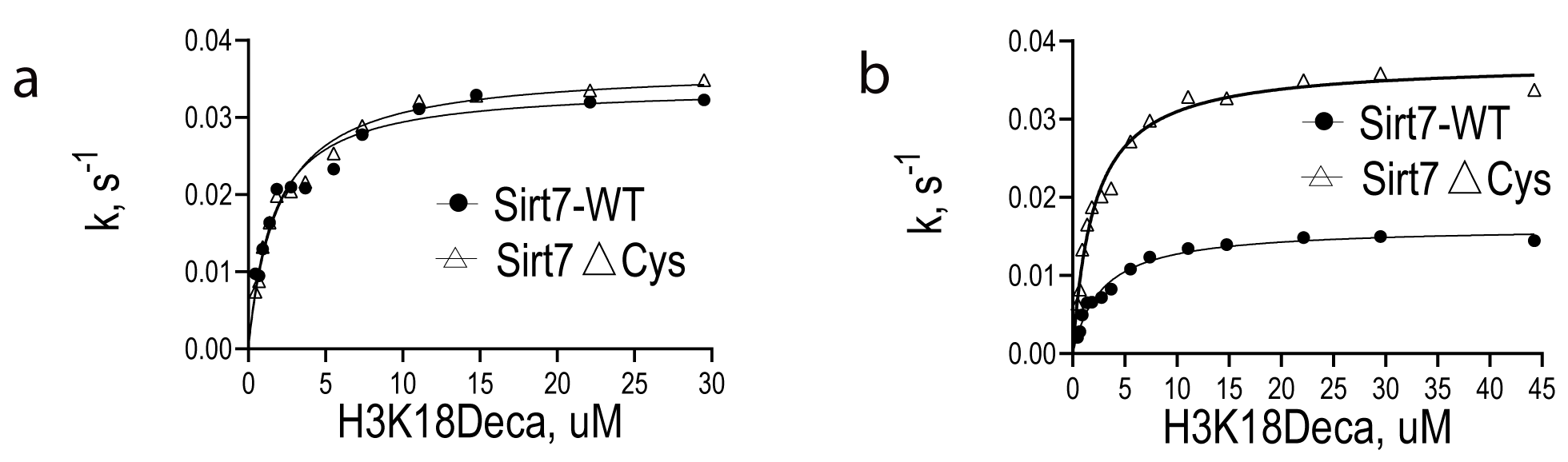

**Supporting figure S2**. Oxidative instability of Sirt7. a. Michaelis Menten kinetic profiles of Sirt7WT and Sirt7ΔCys immediately after purification. b. Michaelis Menten kinetic profiles of flash-frozen aliquots of Sirt7WT and Sirt7ΔCys after 2 month of storage at -80^o^C.

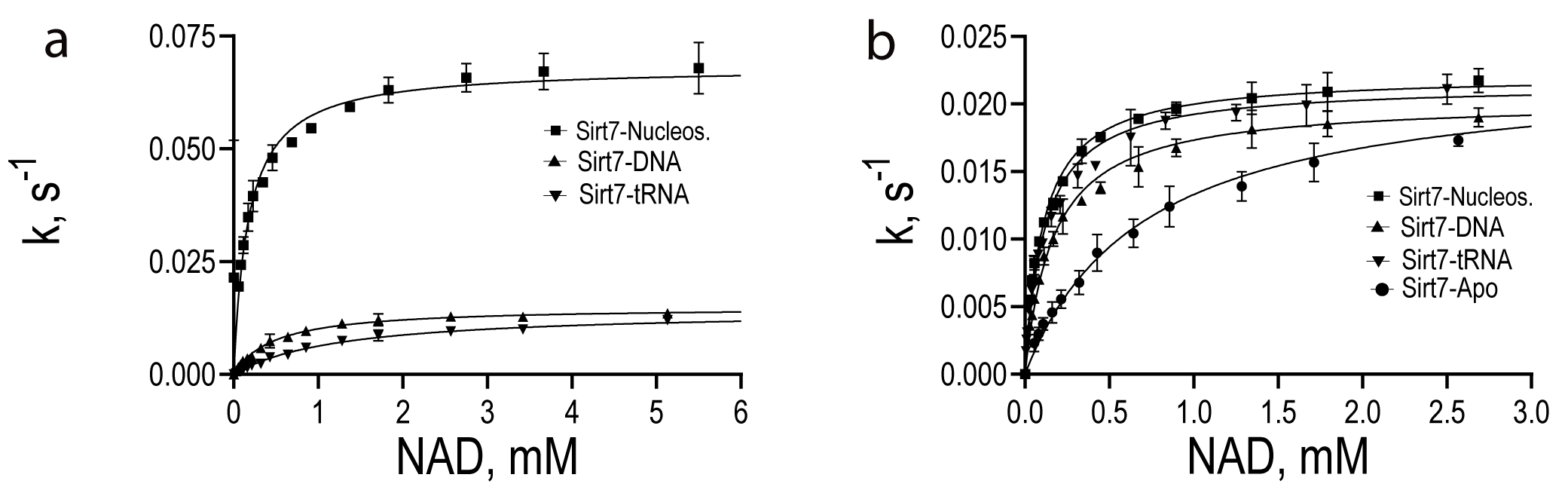
**Supporting figure S3**. Michaelis Menten Sirt7 profiles with variable NAD^+^ substrate and saturating H3K18Ac (a) and H3K18Deca (b) substrates. Corresponding kinetic parameter are summarized in the Table 2, Figure 4.

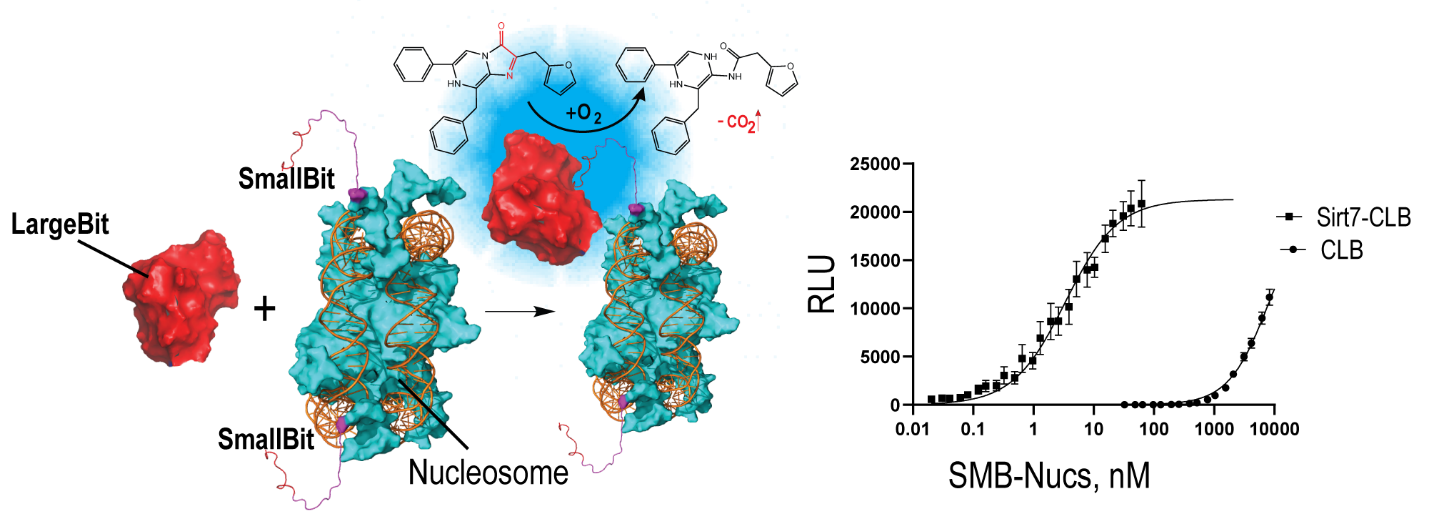

**Supporting figure S4**. Interaction of Sirt7-CLB fusion and CLB domain with SMB-Nucleosomes. Isolated CLB domain does not interact with SMB-nucleosomes up to 1000 nM, and thus doesn’t affect interaction of Sirt7-CLB with SMB-Nucs.
